# Grid Cell Firing Patterns Maintain their Hexagonal Firing Patterns on a Circular Track

**DOI:** 10.1101/2023.09.14.557783

**Authors:** Man Yi Yim, Steven Walton, Kathryn Hedrick

## Abstract

In an open two-dimensional environment, grid cells in the medial entorhinal cortex are known to be active in multiple locations, displaying a striking periodic hexagonal firing pattern covering the entire space. Both modeling and experimental data suggest that such periodic spatial representations may emerge from a continuous attractor network. According to this theory, grid cell activity in any stable 1D environment is a slice through an underlying 2D hexagonal pattern, which is supported by some experimental studies but challenged by others. Grid cells are believed to play a fundamental role in path integration, and so understanding their behavior in various environments is crucial for understanding the flow of information through the entorhinal-hippocampal system. To this end, we analyzed the activity of grid cells when rats traversed a circular track. A previous study involving this data set analyzed individual grid cell activity patterns separately, but we found that individual grid cells do not provide sufficient data for determining the underlying spatial activity pattern. To circumvent this, we compute the population autocorrelation, which pools together population responses from all grid cells within the same module. This novel approach recovers the underlying six-peak hexagonal pattern that was not observable in the individual autocorrelations. We also use the population autocorrelation to infer the spacing and orientation of the population lattice, revealing how the lattice differs across environments. Furthermore, the population autocorrelation of the linearized track reveals that at the level of the population, grid cells have an allocentric code for space. These results are strong support for the attractor network theory for grid cells, and our novel approach can be used to analyze grid cell activity in any undersampled environment.

## INTRODUCTION

Grid cells in the medial entorhinal cortex (MEC) are an important component of the brain’s entorhinal-hippocampal system for spatial memory and navigation. The firing fields of grid cells form a striking hexagonal lattice within open-field environments (Hafting et al., 2005; Rowland et al., 2016). Grid cells are believed to be organized into discrete modules, where the hexagonal lattice of each grid cell within a module has a similar spacing and orientation but may differ in phase (Stensola et al., 2012). Since the MEC is the primary source of spatial information for hippocampal place cells, it is important to understand the behavior of grid cells in order to understand the flow of information through the entorhinal-hippocampal system (Yim et al., 2021).

Grid cells are generally believed to play an important role in path integration, which typically means the brain’s ability to update its internal representation of the animal’s location within an environment using only self-motion cues, such as the animal’s running speed and heading direction (McNaughton et al., 2006; Savelli and Knierim, 2019). However, grid cell activity is also dependent on allocentric cues, such as cue cards placed on the floor or walls of an environment (Fyhn et al., 2007). The manner in which grid cells incorporate allocentric information with the internal neural processing involved in path integration is an active area of research (Knierim et al., 2014; Campbell et al., 2018; Jacob et al., 2019).

The most widely accepted theory regarding grid cell behavior centers on the broad theory of continuous attractor neural networks (CANNs). A CANN is a network of neurons whose population activity vector converges in time to a continuous low-dimensional manifold (Trappenberg, 2002; Wu et al., 2008; Knierim and Zhang, 2012; Poll et al., 2016; Geiger and Hedrick, 2021). According to this theory, the instantaneous population activity of grid cells within a module can be arranged as a hexagonal lattice, and self-motion cues drive translations of this lattice (Burak and Fiete, 2009). Hence, the population activity vector can be described using only the two-dimensional phase of the lattice, resulting in a two-dimensional manifold.

Although it is difficult to directly test the theory of attractor networks, there is experimental evidence that grid cell modules can be considered CANNs. In particular, the spacing and orientation are roughly the same for all grid cells within a module (Stensola et al., 2012; Yoon et al., 2013). Though the hexagonal lattice can expand, contract, rotate, or even become distorted between environments, these changes to the spacing and orientation tend to be coherent for all grid cells within a module (Yoon et al., 2013). Similarly, the relative phase between comodular grid cells tends to be constant across environments, even while the lattice of individual cells may shift (Yoon et al., 2013). Accordingly, there are several computational models for grid cells based on the theory of CANNs (Fuhs and Touretzky, 2006; McNaughton et al., 2006; Guanella et al., 2007; Burak and Fiete, 2009). All such models assume that the population maintains a rigid hexagonal lattice in any environment, and translations of this lattice lead to a hexagonal lattice in the firing patterns of individual grid cells.

In sufficiently large open-field environments, the characteristic hexagonal lattice is easily observable in recordings of individual grid cells. The spatial structure is much less clear, however, when the animal is placed on a track, since the animal’s trajectory undersamples any underlying two-dimensional spatial pattern. In these environments, more sophisticated analytical tools are needed to determine the underlying spatial structure in the grid cell activity patterns. For example, Fourier analysis has revealed that grid cell activity on a linear track is most likely a linear slice of an underlying hexagonal lattice (Yoon et al., 2016). Similarly, approaches from topology have led to the conclusion that grid cell activity on a pinwheel is part of an underlying hexagonal lattice (Gardner et al., 2022). A recent study, however, concluded that grid cells behave differently on a circular track, in which grid cell activity does not appear to be a circular slice of an underlying hexagonal lattice (Jacob et al., 2019). This latter conclusion challenges the CANN theory for grid cells.

Understanding the behavior of grid cells on a circular track is particularly important, as many experimental studies constrain the animal’s path to a circular track (Knierim et al., 2014; Campbell et al., 2018; Jacob et al., 2019; Savelli and Knierim, 2019). Such an environment is particularly useful in studies of path integration, as there is no physical discontinuity that would reset the cellular estimate of the animal’s position.

In this study we use a novel technique to further analyze the data of grid cells recorded on a circular track, presented in Jacob et al. (2019). In particular, we focus on the reported observation that grid cell firing patterns are not a circular slice of a hexagonal lattice (Jacob et al., 2019). This claim was based on two observations. First, they found that the Fourier transform of the linearized activity does not have the characteristic three peaks expected of grid cells on a linear track (Yoon et al., 2016). However, these three peaks are due to the specific geometry of a linear slice of a hexagonal lattice and are not expected given the geometry of a circular track. Second, they observed a low correlation between the grid cell activity on the circular track and the corresponding slice of the grid cell activity in the open arena. However, they did not examine whether the activity on the circular track could be a slice of a hexagonal lattice that differs in spacing, orientation, or phase from that in the open arena. (See Discussion for a further description of this study’s results.)

We use a novel approach to analyze the same data recorded in Jacob et al. (2019) in order to clarify how one should interpret grid cell activity on a circular track. Rather than analyzing individual grid cell activity patterns as was previously done, we compute the 2D population autocorrelation, i.e. the sum of the 2D autocorrelations of individual grid cells within a module. We begin by applying this novel approach to simulated data and exploring its robustness. We then show how the population autocorrelation reveals an underlying hexagonal lattice in the populations of grid cells recorded from two animals. We also use the population autocorrelation to infer the spacing and orientation of the population lattice, revealing how the lattice differs across environments. The population autocorrelation can only reveal an underlying spatial pattern if the firing fields are stable across laps, which was not apparent in the original analysis of individual grid cells (Jacob et al., 2019). We analyze the spatial stability by computing the population autocorrelation on the linearized track, showing that the grid code is an allocentric code at the level of the population. Unlike other methods used to analyze grid cells in undersampled environments (Gardner et al., 2022), our approach can be used when there are a limited number of grid cells and noisy data. Altogether, these results further support the attractor network theory for grid cells and provides a powerful method for analyzing grid cell activity in undersampled environments.

## RESULTS

### Theory: Population autocorrelation reveals the underlying spatial structure

We begin by using numerical simulations to demonstrate the utility of the 2D population autocorrelation in revealing the underlying spatial structure. The model grid cells have idealistic hexagonal firing fields, as shown in Fig. 1A (far left).

**Figure 1.**
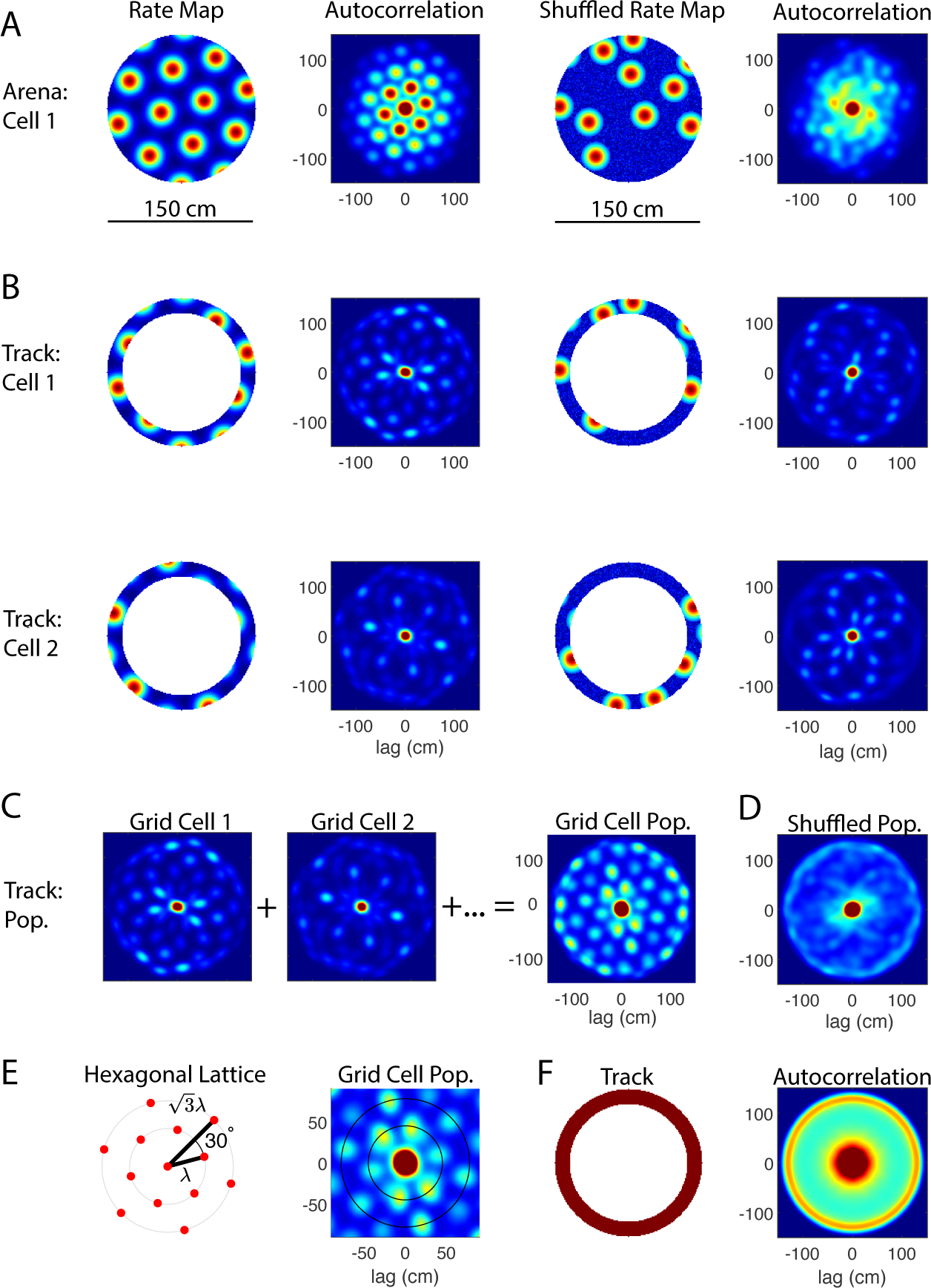
Autocorrelation analysis of simulated rate maps. (A) Simulations of grid cells in an open circular arena (diameter of 150 cm). The 2D autocorrelation of an individual grid cell’s rate map clearly shows the rings of six peaks spaced 60*^◦^* apart that are characteristic of the grid cell hexagonal lattice (left). There is no such spatial structure in the 2D autocorrelation of the corresponding shuffled rate map (right). (B) Two examples of grid cell activity when the simulated rat is constrained to a circular track (outer diameter of 150 cm, width of 15 cm). In simulations, the grid cell activity is a circular slice of the underlying hexagonal lattice (left), where the underlying lattice for Cell 1 is shown in A, and the lattice used for Cell 2 has the same spacing and orientation but a different phase. Because the 2D lattice has been undersampled by the rat’s trajectory, the 2D autocorrelation sometimes shows the characteristic six peaks (Cell 2) and sometimes does not (Cell 1). In general, it is not possible to distinguish between the autocorrelations of the slices of the grid cell rate map from those of the slices of the corresponding shuffled rate map (right). (C) Population autocorrelation of twenty comodular grid cells. The population autocorrelation, or sum of the individual autocorrelations, reveals the signature six-peak hexagonal lattice. (D) Population autocorrelation of the shuffled rate maps. In constrast to (C), the population autocorrelation of the shuffled rate maps has no grid-like spatial structure. (E) Geometry of the hexagonal lattice. The rings passing through the two innermost hexagons have a radius of *λ* and 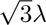, respectively, where *λ* is the grid cell spacing. The second ring is rotated 30*^◦^* from the first. Depending on the size of the environment, the population autocorrelation may reveal multiple six-peaked rings (right, magnified from C). (F) Autocorrelation of the circular track with a uniform firing rate. The circular track contributes a central peak and outer ring to the autocorrelation. The radius of the outer ring corresponds to the diameter of the circular track.

First consider the 2D autocorrelation of individual grid cells. Autocorrelation has been widely applied to reveal the spatial structure of individual grid cells in open-field environments (e.g., Hafting et al. (2005)). In the example shown in Fig. 1A (left), the autocorrelation of the rate map is also hexagonal, with the same spacing and orientation as the rate map itself. Given the finite physical space of the environment, the intensity of the autocorrelation is greatest at the origin and lessens with the radial distance from the origin. In contrast, when the firing fields are shuffled in the same environment, the 2D autocorrelation of the corresponding shuffled rate map generally lacks a hexagonal spatial structure (Fig. 1A (right)).

The effectiveness of this standard approach depends not only on the size of the environment, but also on how thoroughly the animal explores the environment. When the underlying 2D environment is undersampled, such as when the animal is constrained to a circular track, many of the firing fields are “missing” from the rate map. Two examples of this are shown in Fig. 1B (left). In these simulations, the rate map is a perfect, idealistic circular slice of an underlying hexagonal lattice. In the corresponding autocorrelations, however, the characteristic rings of six peaks seen in (A) are now either much less apparent (e.g. Cell 2) or completely absent (e.g. Cell 1). On the other hand, the 2D autocorrelation of the shuffled rate maps can sometimes resemble the ring of six peaks (Fig. 1B (right)). Even in this idealistic setting, the 2D autocorrelation of individual rate maps cannot be used to determine whether the activity is a circular slice of a hexagonal lattice because they are indistinguishable from the autocorrelations of the shuffled rate maps.

We circumvent this limitation by computing the population autocorrelation, defined as the sum of the autocorrelations of individual rate maps of all grid cells within a module. For the example shown in Fig. 1C, the population consists of twenty grid cells, each with a hexagonal lattice that has the same spacing and orientation but differs in phase, as expected of idealistic grid cells within the same module (Stensola et al., 2012). The resulting population autocorrelation (C, right) shows a clear hexagonal lattice, with the same interior peaks as found in the arena (A) but with a different distribution of intensities. In contrast, no spatial structure emerges in the population autocorrelation for the corresponding shuffled rate maps (Fig. 1D).

This approach is effective because the autocorrelation is independent of phase. In these simulations, the lattices of the comodular grid cells have exactly the same spacing and orientation, differing only in phase. Thus, the individual autocorrelations would be identical in sufficiently large and well-sampled environments. In undersampled environments such as a circular track, each autocorrelation is missing different peaks because the intersection of the underlying hexagonal lattice and the circular track depends on the phase of the lattice. Hence, the full hexagonal lattice emerges when the individual autocorrelations are combined, and the intensity of each peak depends in part on how many individual grid cells show that peak in its autocorrelation. The opposite effect occurs in the shuffled population. While some individual autocorrelations may appear hexagonal by chance, there is not a consistent spacing and orientation among the shuffled rate maps. This results in a population autocorrelation that shows no spatial structure, allowing one to clearly distinguish the circular slice of a hexagonal lattice from that of a shuffled rate map.

Fig. 1E shows further detail of the geometrical structure of the hexagonal lattice expected in the population autocorrelation. The ideal autocorrelation consists of concentric rings, each containing six peaks uniformly spaced 60*^◦^* apart. The radius of the innermost ring is the spacing of the grid cell module (*λ*). The second ring is rotated 30*^◦^* with respect to the first and has radius 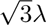. Note that in some cases, the intensity of the peaks in the second ring is greater than those in the first ring. Thus, the most prominent ring may not be the innermost ring, particularly when analyzing noisy experimental data.

Independent of the firing fields, the geometry of the circular track contributes a large circular peak and outer ring to the 2D autocorrelation (Fig. 1F). The central peak is typical of any autocorrelation, and here it reflects the size of individual firing fields. The outer ring corresponds to the diameter of the track and reflects the fact that the firing fields are constrained to the circular track. Consequently, one cannot draw any conclusions about the underlying spatial structure using the central peak or the outer ring of peaks in the autocorrelation. (Note that in C and D, the outer ring of peaks is clearly visible, and the central peak is slightly larger than in the individual autocorrelations shown in B.)

### Theory: Results are robust to more realistic conditions

To verify that the results presented in Fig. 1 are robust, we next show that the population autocorrelation remains effective given conditions that more closely resemble experimental data. Specifically, we test the robustness of the results for three parameters: the spacing of the lattice, the thickness of the track, and variability in the spacing and orientation of grid cells within a module. Fig. 2 shows results from one representative example of each parameter set.

**Figure 2.**
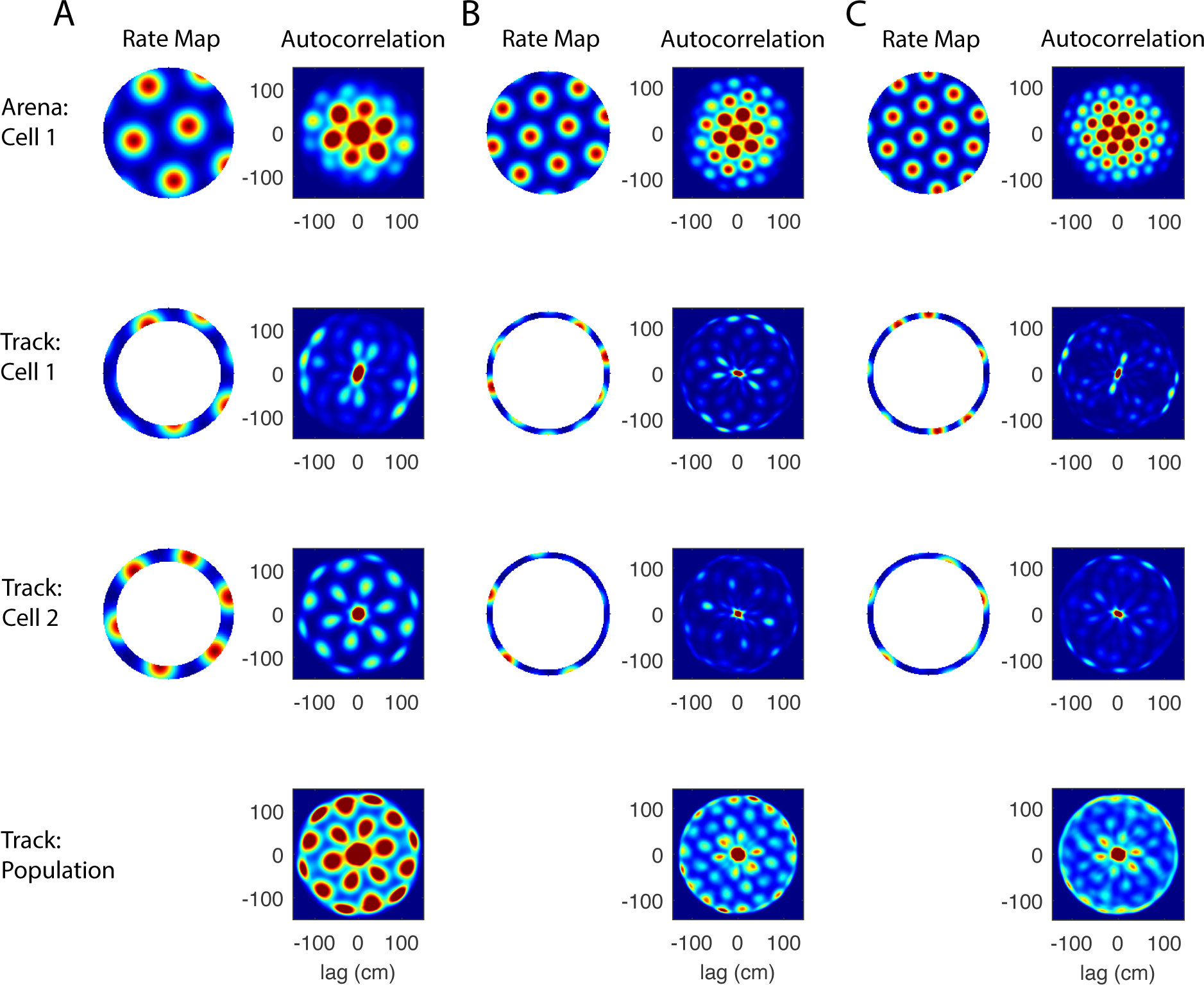
Robustness of the population autocorrelation analysis. (A) Robustness to larger grid spacing. The population autocorrelation (bottom) still reveals the underlying hexagonal lattice when the grid spacing is increased from 45 cm (Fig. 1) to 65 cm here. The innermost six-peaked ring is clearly visible, but the second ring is now partly obscured by the effects of the annulus (Fig. 1F). (B) Robustness to a thinner track. The grid parameters are the same as used in Fig. 1, except here the track is 7 cm wide instead of 15 cm wide. This results in a greater undersampling problem, but the six-peaked rings are still revealed by the population autocorrelation (bottom). This thinner track may be a more realistic representation of a rat’s trajectory on an annulus, since rats often do not explore the entire width of the track. (C) Robustness to variability in the spacing and orientation. In addition to the thinner track, the spacings and orientations of the twenty grid cells were varied in accordance with the natural variability observed in experimental recordings. The six-peaked population autocorrelation remains clearly visible (bottom).

The spacing and size of grid fields increase from dorsal to ventral MEC, and the observed spacing is within the range of about 30 to 150 cm (Hafting et al., 2005; Stensola et al., 2012). The population autocorrelation is most effective when applied to the smaller grid fields located in the dorsocaudal MEC, as a smaller spacing leads to more distinct grid fields on the track. To test our approach, we increase the lattice spacing from 45 cm (used to generate Fig. 1) to 65 cm (Fig. 2A). The hexagonal lattice is clearly visible in the population autocorrelation (Fig. 2A, bottom), and the larger field size and spacing in the autocorrelation (compared to Fig. 1) reflects the larger field size and spacing in the rate map. The two innermost rings of six peaks are both apparent, though the second ring of six peaks is somewhat obscured by the effects of the track (see Fig. 1F). In this example, the second ring has a radius of about 113 cm, and the inner diameter of the track is 120 cm. Hence, the six peaks of the second ring are still apparent but overlap with the smaller peaks that are due to the track itself. The inner diameter of the track is an upper bound on the lattice spacing for which the method may be used.

Second, we show that the population autocorrelation is robust to the thickness of the track (Fig. 2B). The 2D autocorrelation of an individual grid cell will generally show a more complete hexagonal lattice when the animal explores a larger proportion of the underlying 2D space. We find that the population autocorrelation still reveals a clear hexagonal lattice when the track width is decreased from 15 cm (used to generate Fig. 1) to 7 cm, though the inner ring of six peaks becomes a bit distorted in this example (Fig. 2B, bottom). Note that the width of the track used in simulations best represents the width of the animal’s trajectory, not necessarily the width of the physical track, and 7 cm is a reasonable lower bound for the trajectory width.

Finally, in addition to using the thinner track, we varied the spacing and orientation of each individual grid lattice by drawing these parameters from the normal distributions 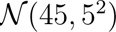(cm) and 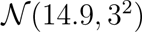 (*^◦^*), respectively. This variability is in accordance with the natural variability of the grid lattice observed in recordings. We again observed a clear hexagonal lattice structure in the population autocorrelation, though the second ring of six peaks is not as clear for this example (Fig. 2C).

Altogether, these simulations indicate that the 2D population autocorrelation is an effective method for determining whether the activity on a circular track is a slice of underlying hexagonal lattice. The resulting hexagonal lattice becomes more clear given a smaller grid spacing and wider animal trajectory. Notably, it requires only a small number of comodular grid cells (20 were used for the simulations presented here), and it is robust to the variability in spacing and orientation expected in experimental data. Next, we apply this approach to experimental data to determine whether a population of grid cells maintains their hexagonal firing patterns on a circular track.

### Autocorrelations of individual grid cells do not reveal a consistent spatial structure on the circular track

We apply our theory to data recorded from rat grid cells (Jacob et al., 2019). We analyze a total of 33 grid cells from two rats (18 in Rat 1 and 15 in Rat 2) recorded in three environments: an open circular arena of diameter 150 cm (*E*_1_) and a circular track of outer diameter 150 cm and width 15 cm in light (*E*_2_) and dark (*E*_3_) conditions. The open arena had a uniformly black wall with one polarizing white cue card attached, which was removed only in the dark condition. The track was created from the open arena by inserting an opaque circular wall into the arena, and the animal was briefly removed and disoriented between recording sessions (Jacob et al., 2019). This data set is appropriate for testing our theory because the data in the open arena can be used to determine which cells fire in the hexagonal lattice pattern that is characteristic of grid cells. Since the same cells were also recorded on the circular track, we can then use our method to determine whether these grid cells maintain an underlying 2D hexagonal lattice even when the animal is constrained to a circular track. If so, the method can be further used to determine if there are changes such as an expansion, contraction, or rotation to the lattice.

Fig. 3 shows the trajectory, rate map and autocorrelation of four example grid cells. All four cells are classified as grid cells due to the clear hexagonal structure in the open arena (*E*_1_), as indicated by the six prominent peaks spaced 60*^◦^* apart in the autocorrelation. As expected from our numerical simulations, the autocorrelation in the circular track environments (*E*_2_ and *E*_3_) do not show a clear, consistent spatial structure. Of the two example cells in Rat 1, Cell 4 displays a noisy six-peaked pattern in both the light and dark conditions (A), whereas Cell 9 reveals a noisy six-peakedness in the light condition only (B). Cell 12 in Rat 2 does not show a six-peaked pattern in either condition (C). See Supp. Figs. S1-S3 and S8-S10 for autocorrelations of all grid cells in Rats 1 and 2 in all three environments.

**Figure 3.**
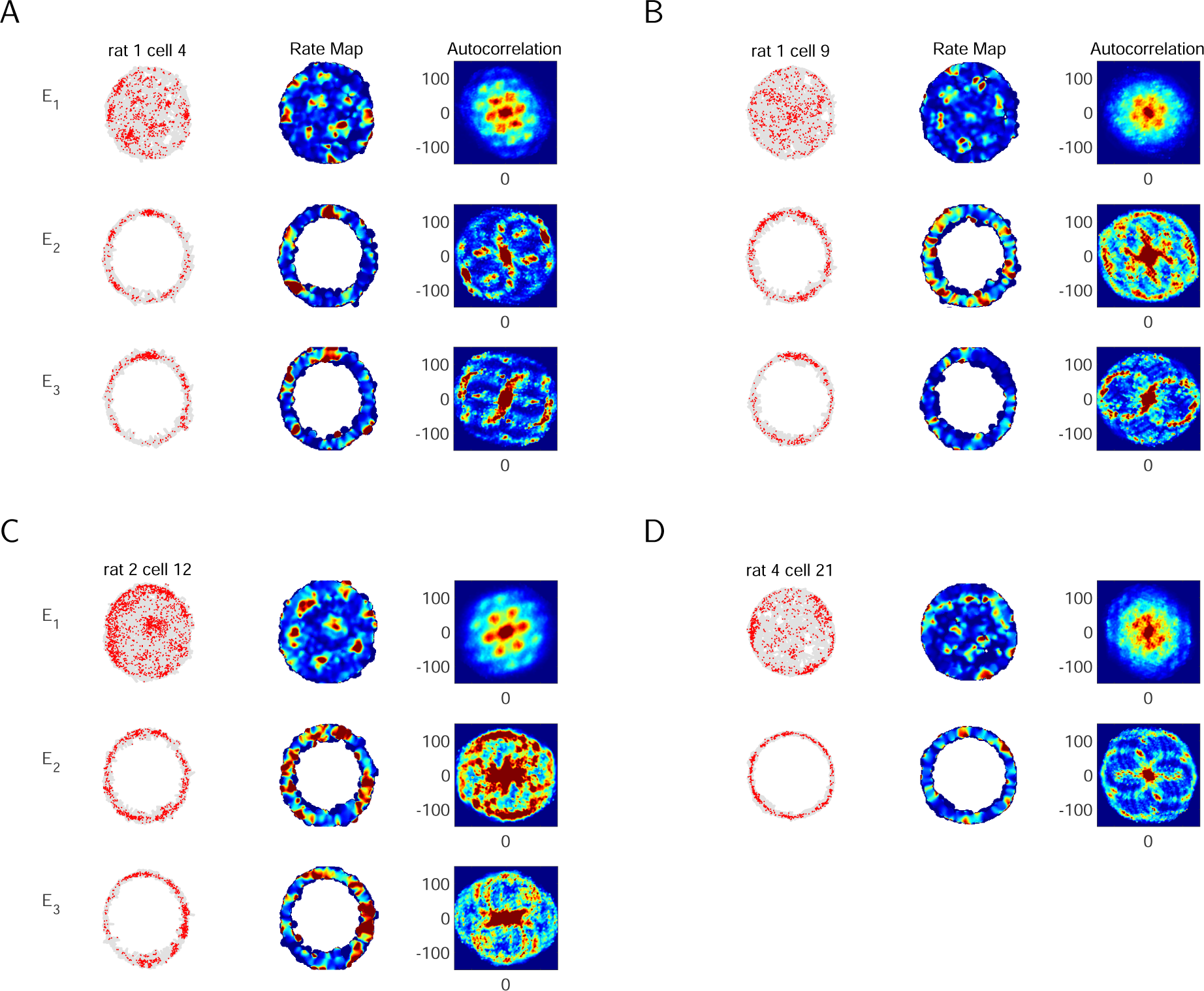
Examples of single-cell analysis of experimental data from an open circular arena and a circular track. The 2D autocorrelation of individual grid cells reliably shows a 2D hexagonal lattice in the open-field environment (*E*_1_), which is the standard approach used to classify grid cells. However, as expected from numerical simulations, the same 2D autocorrelation applied to data from individual grid cells does not reliably reveal an underlying spatial structure on the circular track. (A) An example cell in Rat 1. In the plots on the left, each red dot shows the location of the rat when this grid cell fired an action potential, and the gray shows the rat’s trajectory. In the open arena (*E*_1_, top), the 2D autocorrelation (right) of the rate map (middle) for this individual cell clearly shows the six-peaked ring of the innermost hexagon. However, there is not a clear spatial structure in the autocorellation from the circular track in the light condition (*E*_2_, middle) or dark condition (*E*_3_, bottom). (B-D) Same as A, but for another example cell in Rat 1 (B), an example cell in Rat 2 (C), and an example grid cell in Rat 4 (D). (Data in *E*_3_ for Cell 21 of Rat 4 was not available.)

Of the published data, only data from Rats 1 and 2 contained multiple cells that were recorded in all three environments and that we clearly classified as grid cells in the open arena. However, we include one cell from Rat 4 as an additional example of a grid cell that displays a clear hexagonal pattern on the circular track (D). However, as this can happen by chance, one cannot use this example to draw general conclusions about the behavior of grid cells on a circular track.

### Population autocorrelation reveals an underlying hexagonal lattice in Rat 1

Based on our numerical simulations, the population autocorrelation should reveal an underlying hexagonal lattice on the circular track if and only if (1) grid cell firing patterns are a circular slice of an underlying 2D hexagonal lattice, and (2) the data contains a sufficient number of grid cells with a similar orientation and spacing, as would be expected of grid cells within the same module. Regarding (2), it is reasonable to consider the grid cells recorded from Rat 1 as being comodular because the population autocorrelation from the open arena has a clear hexagonal structure with a higher gridness score than in any of the individual autocorrelations (Fig. 4A). (See Supp. Fig. S1 for all individual autocorrelations in *E*_1_.) This can only occur when the individual autocorrelations align well with one another. We show this explicitly by inferring the orientation and spacing of each individual grid cell in the open arena. As shown in Fig. 4E, the orientations of all grid cells cluster around three angles 60*^◦^* apart, while the spacing of most grid cells clusters around 45 cm.

**Figure 4.**
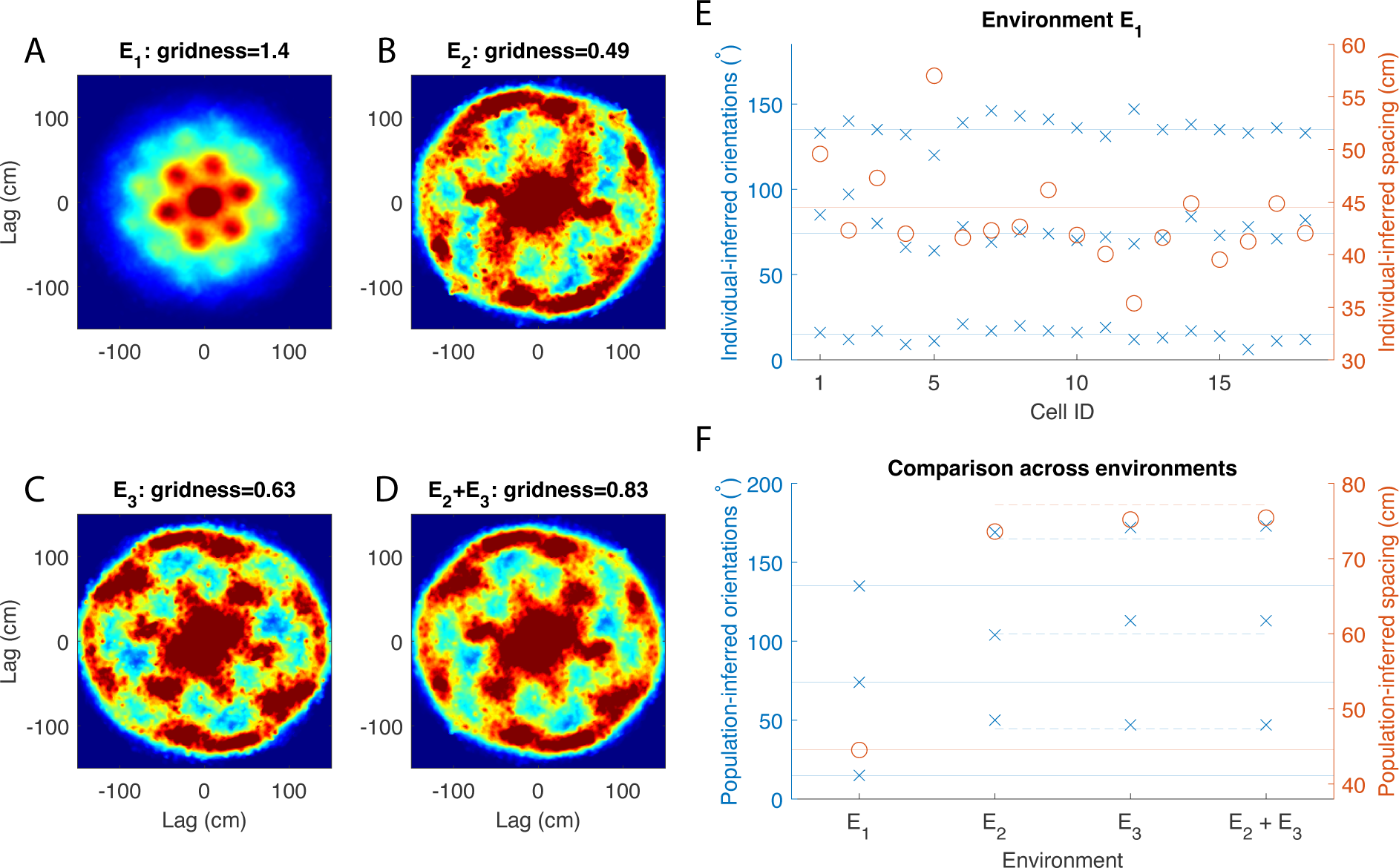
Population analysis of grid cells in Rat 1. (A) Population autocorrelation for the open arena (Environment *E*_1_). While the autocorrelations of the individual grid cells show a hexagonal lattice, the population autocorrelation shows a more prominent hexagonal pattern with a higher gridness score. This indicates that the grid spacing and orientation are consistent across cells. (B-C) Population autocorrelation for the circular track in the light (B) and dark (C) conditions. In both cases, the population autocorrelation recovers the six-peak hexagonal signature of grid cells that is absent in the individual autocorrelations (Supp. Fig. S2-S3). (D) Sum of the population autocorrelation in the light condition (shown in B) with that in the dark condition (shown in C). Taking the sum of all 16 autocorrelations from the circular track amplifies the six-peak hexagonal signature, as indicated by a higher gridness score. (E) Inferred spacing orientations from the open arena. The solid lines indicate the inferred spacing (red) and orientation angles (blue) of the population autocorrelation. The inferred parameters from the 18 individual grid cells cluster around the parameters from the population autocorrelation. (F) Inferred spacing and orientations from each environment, computed from the population auto c<u>o</u>rrelations. Solid lines show the parameters from the open arena (same as in E). The dashed orange line shows 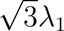, where *λ*_1_ is the population spacing from the open arena. The dashed blue line shows a 30*^◦^* rotation between the orientations in open arena. Results indicate the grid spacing and orientation are the same across all environments, and the population autocorrelation on the circular track shows the second ring of six peaks (see Fig. 1E).

Assuming the grid cells recorded in Rat 1 are indeed comodular, the population autocorrelation can be used as a test for the hypothesis (1) for Rat 1. Fig. 4B and C show the population autorocorrelation, or sum of the individual autocorrelations of the 18 grid cells, in the light and dark conditions, respectively. A clear ring of six peaks spaced approximately 60*^◦^* apart emerges with a high gridness score, particularly in the dark condition. Notably, this ring of six peaks in both conditions is much more prominent than in any of the autocorrelations of individual grid cells in either environment (shown in Supp. Fig. S2-S3). Therefore, the population autocorrelation provides strong evidence that this population of grid cells maintains an underlying hexagonal lattice in the circular track environments.

Environments *E*_2_ and *E*_3_ are the same circular track under different conditions (light and dark, respectively), and we combined them together by summing the two autocorrelations for each individual grid cell (shown in Supp. Fig. S4). If the underlying hexagonal lattice is preserved across the two conditions, then the combined autocorrelation effectively incorporates more data for each grid cell. We computed the population autocorrelation for the combined environments by taking the sum of these 18 combined autocorrelations. The hexagonal pattern is reinforced in the population autocorrelation of the combined environments (Fig. 4D), producing an even higher gridness score than in the two environments individually (Fig. 4B-C). This indicates that the grid spacings and orientations are consistent across the light and dark conditions on the circular track. The individual grid cell autocorrelations in the combined environments typically do not show a clear hexagonal structure. One interesting exception to this is Cell 8, which has the characteristic evenly-spaced six peaks in the combined environments (Supp. Fig. S4), despite showing only four peaks in Environment *E*_2_ (Supp. Fig. S2) and two peaks in Environment *E*_3_ (Supp. Fig. S3). This could be explained by a phase shift between the light and dark conditions, resulting in different firing fields aligning with the track. Cell 8 exemplifies the basic idea behind the theory presented here, though in general several autocorrelations are required in the summation to complete the hexagonal pattern (i.e. by taking the population autocorrelation).

It is important to emphasize that the population autocorrelation in all three cases (*E*_2_, *E*_3_, and *E*_2_ + *E*_3_) shows a much clearer hexagonal pattern than in any of the individual grid cell autocorrelations. The spatial structure is a property of the population and is not due to a few dominant grid cells. To verify this, we removed the three grid cells with the clearest hexagonal structure in the combined environments (Cells 14, 8 and 16 in Supp. Fig. S4) and recomputed the population autocorrelation. The hexagonal structure in the resulting population autocorrelation is robust to removing these cells, and the gridness score remains high (Supp. Fig. S5, bottom) despite the lack of hexagonal structure in any of the remaining autocorrelations for individual grid cells.

### Grid spacing and orientation are consistent across environments in Rat 1

Due to its prominent ring of six peaks, one can use the population autocorrelation to infer the grid spacing and orientation of the hexagonal lattice in each environment (see Methods for details). In the open arena (Environment *E*_1_), the population autocorrelation has a grid spacing of 44.5 cm and grid orientation angles of 15*^◦^,* 74*^◦^* and 135*^◦^*. As expected, the orientation angles are spaced about 60*^◦^* apart. Because autocorrelation of individual grid cells also shows a hexagonal pattern in the open arena, we compare the inferred population parameters to those of each individual grid cell (Fig. 4E). The individual grid spacings range from 35.4 to 57 cm, with most values clustering near the inferred population spacing. The individual grid orientations also cluster tightly around the population orientation angles. This is further evidence that all grid cells in Rat 1 may be part of the same module, and the population autocorrelation provides an accurate measure of the grid spacing and orientation of a module.

We next infer the population grid spacing and orientation in the circular track environments and compare the grid parameters across environments (Fig. 4F). The grid spacing and orientation on the circular track are roughly the same in both the light and dark conditions, which is why the population autocorrelation shows a clearer hexagonal pattern in the combined *E*_2_ + *E*_3_ environment. Interestingly, the inferred spacings on the circular track closely approximate 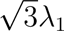 (F, orange dahsed line), where *λ*_1_ is the population grid spacing from the open arena. Furthermore, the inferred orientations on the circular track closely approximate the midpoints between the orientations from the open arena (F, blue dashed lines).

It is possible that the hexagonal lattice expands by a factor of 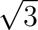 and rotates 30*^◦^* on the circular track compared to the open arena. However, based on the geometry of a hexagonal lattice, the most likely explanation is that the grid spacing and orientation are the same in all environments, but the population autocorrelation in the open arena shows the innermost ring of six peaks, while the population autocorrelation in the circular track environments shows the second ring of six peaks. As illustrated in Fig. 1E, this would explain the change in both the spacing and orientation. There is a large central peak in the population autocorrelations of the circular track environments (Fig. 4B-D), which appears to obscure the innermost ring in the hexagonal pattern.

### Population autocorrelation does not reveal a spatial structure when applied to all grid cells in Rat 2

We next performed the same analysis on the population of 15 grid cells recorded in Rat 2. The autocorrelations of each individual grid cell are shown in Supp. Fig. S8-S11, and the population autocorrelation in each environment is shown in Fig. 5A-B. The population autocorrelation shows a clear hexagonal lattice in the open arena (A). Unlike in Rat 1, however, there is no discernible spatial structure in the population autocorrelation for the circular track environment in the light or dark condition (B).

**Figure 5.**
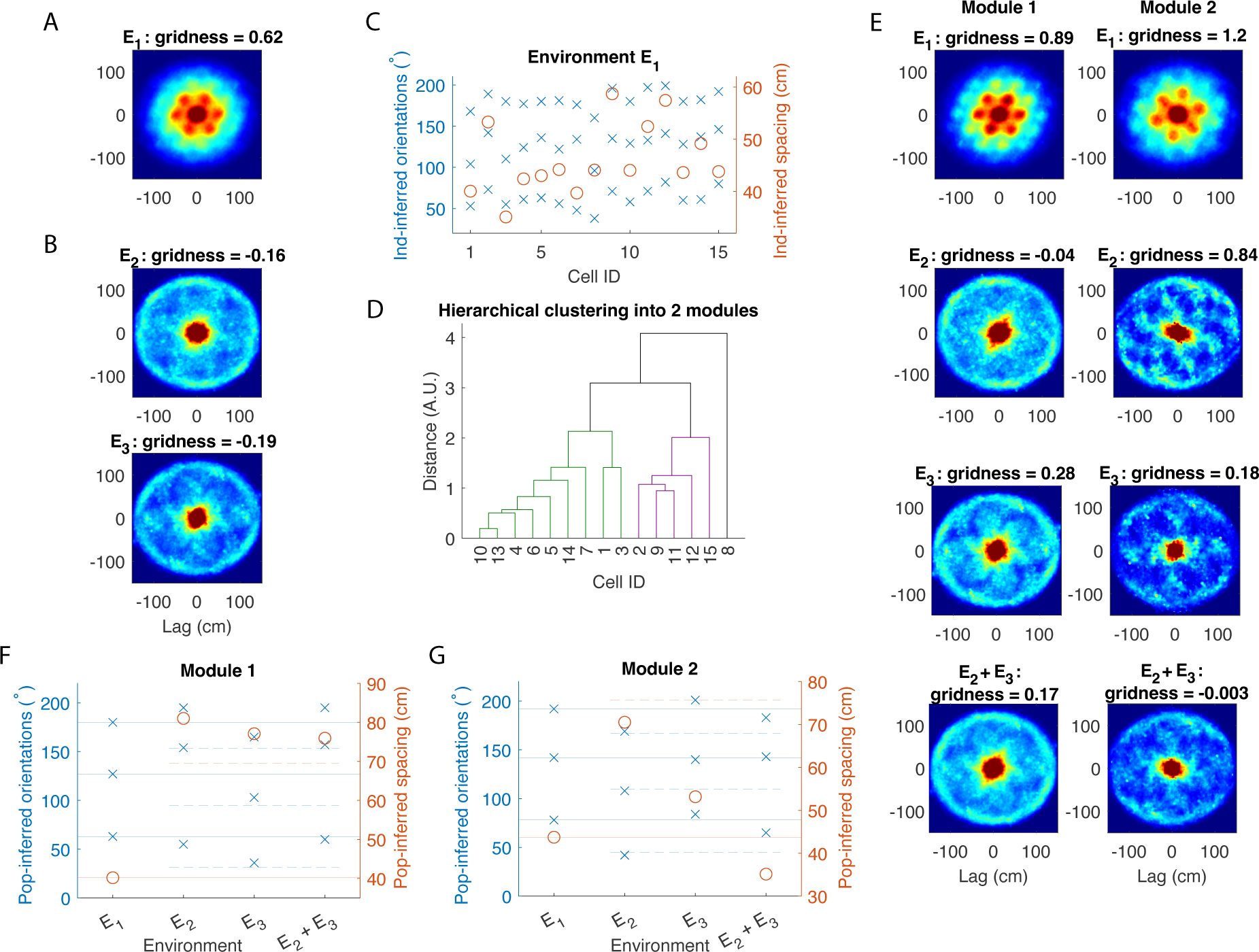
Population analysis of grid cells in Rat 2. (A) Population autocorrelation for the open arena (Environment *E*_1_). The population autocorrelation shows a clear hexagonal lattice, but the gridness score is lower than that of several of the individual grid cell’s autocorrelations (shown in Supp. Fig. S8). This indicates that the firing patterns of individual grid cells do not all align well with one another. (B) Population autocorrelation for the circular track in the light (B) and dark (C) conditions. There is no spatial structure apparent in either condition. (C) Inferred spacing and orientation of each individual autocorrelation for the open arena (Supp. Fig. S8). Unlike in Rat 1, the grid parameters do not cluster around the corresponding parameters of the population autocorrelation. (D) Hierarchical clustering applied to the 15 grid cells based on their inferred spacing and inferred orientations. The grid cells are classified into two modules, where Module 1 has 9 cells and Module 2 has 5 cells. One outlier is discarded in this analysis. (E) Population autocorrelation in each environment for each module. The hexagonal lattice becomes more prominent in the open arena (top row). The results on the circular track are mixed, with a hexagonal structure appearing most clearly in *E*_3_ of Module 1 and *E*_2_ of Module 2. (F-G) Inferred orientations and spacing from the population autocorrelation in each environment for Module 1 (F) and Module 2 (G). The solid lines indicate the orientation angles (blue) and spacing (orange) from the open arena. For the two circular track environments with a clear hexagonal lattice (Module 1 *E*_3_ and Module 2 *E*_2_), the orientations are midway between the orientations from the open arena (dashed blue lines), and the spacing is close to 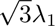 (dashed orange line), where *λ*_1_ is the spacing from the open arena. This indicates that the population autocorrelation again detects the second ring of six peaks, and the grid orientation and spacing is consistent between environments.

In this case, the lack of spatial structure in the population autocorrelation cannot be used as evidence for or against the hypothesis that grid cell activity is a slice of an underlying hexagonal lattice. To understand why, first note that even for the open arena, the hexagonal lattice in the population autocorrelation (Fig. 5A) has a much lower gridness score than that in Rat 1 (Fig. 4A), and it is lower than several of the gridness scores for the individual grid cells in Rat 2 (Supp. Fig. S8). This indicates that the firing fields of the individual autocorrelations in the open arena do not align as well for Rat 2 as they did for Rat 1. Indeed, the individual autocorrelations shown in Supp. Fig. S8 appear to vary widely in orientation and spacing. As shown in Fig. 5C, the inferred orientations and spacings of the individual autocorrelations do not cluster around the corresponding values from the population autocorrelation as they did in Rat 1. Therefore, the population autocorrelation would not uncover a hexagonal pattern even if the hypothesis were true, for the individual autocorrelations would effectively obscure one another in the summation rather than completing one unified pattern.

### An underlying hexagonal lattice emerges when grid cells in Rat 2 are divided into two modules

According to our theory, the population autocorrelation should reveal an underlying hexagonal lattice if the summation is taken over grid cells in the same module (or at least with a similar spacing and orientation). To further analyze the grid cell activity in Rat 2, we applied hierarchical clustering to separate the 15 grid cells into two modules, using the distance between the vectors consisting of each grid cell’s spacing and three orientation angles inferred from the open arena. As shown in the dendrogram in Fig. 5D, the grid cells can be split into two modules, with nine cells in Module 1 and five cells in Module 2. One outlier cell is discarded from this analysis.

Fig. 5E shows the population autocorrelation applied to each module in each environment. As expected, the population autocorrelation for the open arena has a much clearer hexagonal structure for each module (E, top row) than for all 15 grid cells combined (A). The results in the circular track environments are mixed. For Module 1, there is a clear hexagonal structure with a high gridness score in the circular track environment in the dark condition (*E*_3_), but not in the light condition (*E*_2_). For Module 2, there is a clear hexagonal structure in the light condition and a weaker hexagonal structure in the dark condition, but no structure in the combined *E*_2_ + *E*_3_ environment.

### Grid spacing and orientation also appear consistent in Rat 2

As done for Rat 1, we use the population autocorrelation to compare the grid spacing and orientation of the underlying hexagonal lattice in the circular track environments to that in the open arena (Fig. 5F-G). We focus on Module 1 Environment *E*_3_ and Module 2 Environment *E*_2_, as those are the two circular track environments in which the hexagonal lattice is the most apparent. In both cases, the orientation angles closely approximate the midway point between th<u>e</u> orientation angles from the open arena. Furthermore, the spacing is a fair approximation to 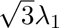, where *λ*_1_ is the spacing of the population autocorrelation for the open arena. Thus, while the results are not as clear as in Rat 1, it is reasonable to conclude that the grid spacing and orientation are again consistent across environments, and the population autocorrelation in the circular track environments detects the second ring of six peaks.

### Gridness scores of the population autorocrrelations are statistically significant

It is possible that firing fields which do not arise from an underlying hexagonal lattice may nonetheless lead to an autocorrelation that appears hexagonal. To test the significance of our results, we generated 1000 controls for each grid cell in each environment. Each control has the same non-spatial statistics as the original data, including the number and size of individual firing fields. Specifically, the controls were generated by shuffling the firing fields of the original rate maps, placing each firing field in a random location without any two fields overlapping. (See Methods for more details.) Supp. Fig. S6 shows examples of controls for the open arena, and Supp. Fig. S7 shows examples for the circular track environments. We then computed the population autocorrelation for each set of controls, comparing the gridness score for the actual data to those of the 1000 controls in each environment (Fig. 6).

**Figure 6.**
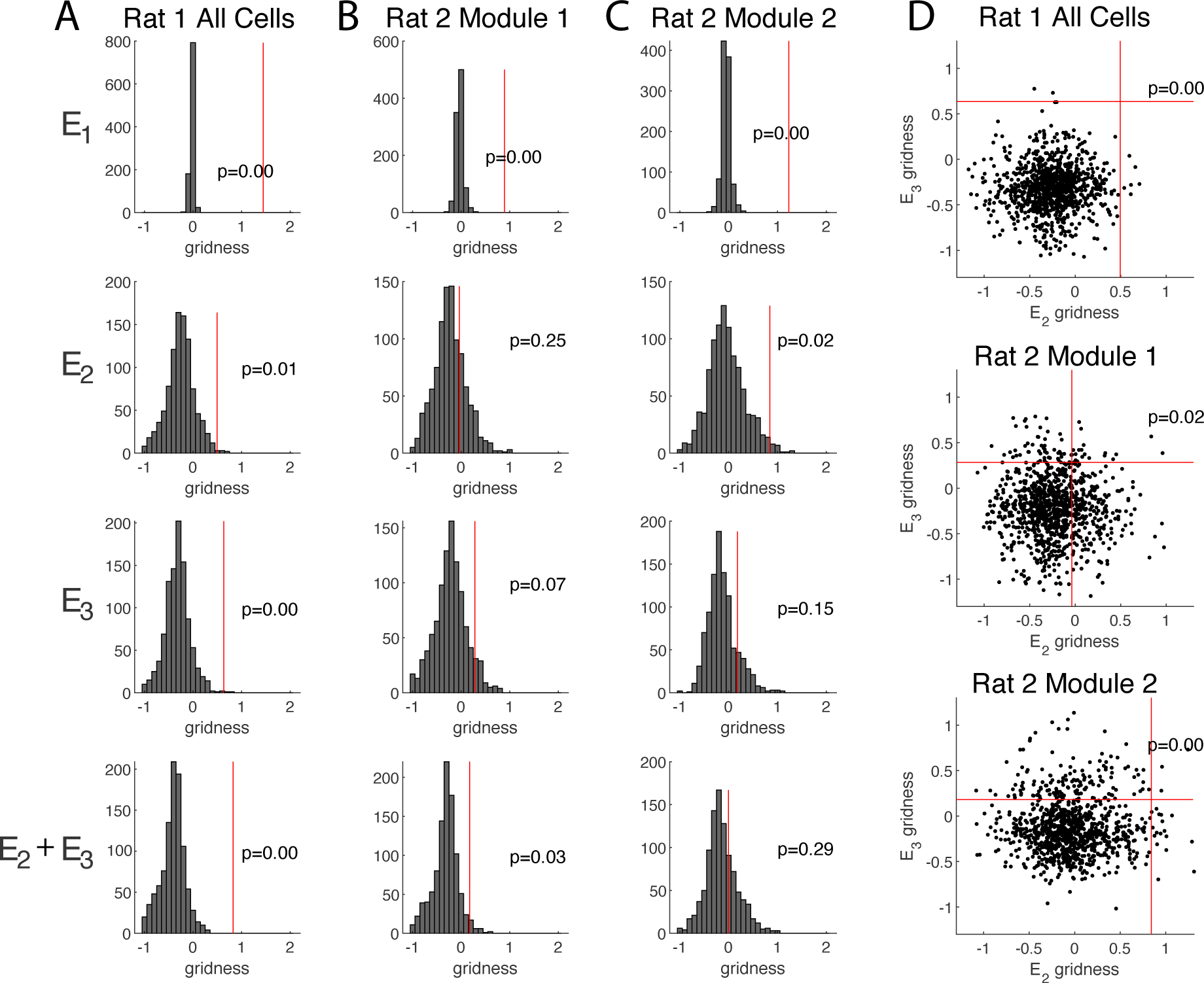
Significance of the hexagonal structure in the population autocorrelation. (A) The vertical red line shows the gridness score of the population autocorrelation of the true data from Rat 1 (Fig. 4), which is compared to the histogram of gridness scores of the population autocorrelations of 1000 shuffle controls (e.g. Supp. Fig. S7). The gridness score is significant in all environments for Rat 1. Indeed, only a few shuffles obtain as high of a gridness score as the true data in *E*_2_ and in *E*_3_, and the true data has a much higher gridness score than all 1000 shuffles in the combined *E*_2_ + *E*_3_ environment. (B-C) Same as (a) but for Rat 2, Module 1 (B) and Module 2 (C). The gridness score is significant in the open field environment for both modules. On the circular track, the gridness score is only significant for the combined *E*_2_ + *E*_3_ environment for Module 1 and the light condition (*E*_2_) for Module 2. (D) Joint distribution of gridness scores from both conditions on the circular track (*E*_2_ and *E*_3_). When compared to this joint distribution, the pair of gridness scores is significant for Rat 1 and for both modules in Rat 2.

For Rat 1, we find that the gridness score of the population autocorrelation of the data is statistically significant in all environments (*E*_1_: *p* = 0.00*, E*_2_: *p* = 0.01*, E*_3_: *p* = 0.00*, E*_2_ + *E*_3_: *p* = 0.00), further supporting the conclusion that in Rat 1, grid cells maintain an underlying hexagonal lattice in the circular track environments.

For Rat 2, we conducted the same test for significance for each module (Fig. 6B-C). As expected, the gridness scores in the open arena are significant for each module. In the circular track environments, only the combined *E*_2_ + *E*_3_ environment for Module 1 and the light condition (*E*_2_) for Module 2 have significant gridness scores (*p* = 0.03 and *p* = 0.02, respectively). As a further test, for each module we compared the pair of gridness scores in *E*_2_ and *E*_3_ to the joint distribution of gridness scores in the light and dark conditions (Fig. 6D). We found that the pair of gridness scores is significant for both Module 1 and Module 2 (*p* = 0.02 and *p* = 0.00, respectively).

The weaker results in Rat 2 compared to Rat 1 can be explained by the relatively few grid cells in each module of Rat 2. Even so, the appearance of a hexagonal lattice when the cells are split into modules, particularly in *E*_3_ of Module 1 and *E*_2_ of Module 2, and the significance of the gridness score in some of the environments provide additional evidence that the grid cells maintain their 2D hexagonal lattice in the circular track environments.

### Linearization of circular track activity supports an allocentric code in grid cells

Lap-by-lap spatial stability in the grid cell firing patterns is a necessary condition for the population autocorrelation to reveal any underlying spatial structure. This is because the autocorrelations are computed using the average firing rates over an entire session, and large shifts in the firing fields over time would obscure this structure. In the original analysis of this data set, the authors concluded that most individual grid cells do not have an allocentric code for space, meaning their firing fields were not stable over multiple laps (Jacob et al., 2019). Here, we determine if the grid code is allocentric on the level of the population by computing the population autocorrelation of the activity on the 1D linearized track, in which the location of each spike is computed as the total angular distance traveled from the animal’s starting location. Note that multiples of the track circumference (471 cm) correspond to the animal having traveled a complete lap around the track and returning to its starting position. Thus, for a grid cell with stable firing fields across multiple laps, the autocorrelation of its average rate along the track would have a peak at the value of the circumference of the track.

The results are shown in Fig. 7. For individual cells, there is no clear spatial structure in the 1D autocorrelation of the linearized firing rates (A-B). Indeed, in most cases the peaks in the autocorrelation cannot be distinguished from the random fluctuations observed in the control. In contrast, the population autocorrelation in all four cases (light and dark conditions of the circular track for Rats 1 and 2) shows a clear peak that coincides with the circumference of the track (C-D). This indicates that at the population level, the spatial locations of the firing fields are somewhat stable and do not have a consistent drift. Thus, the population autocorrelation reveals an allocentric code for space that is not apparent in the autocorrelations of individual cells.

**Figure 7.**
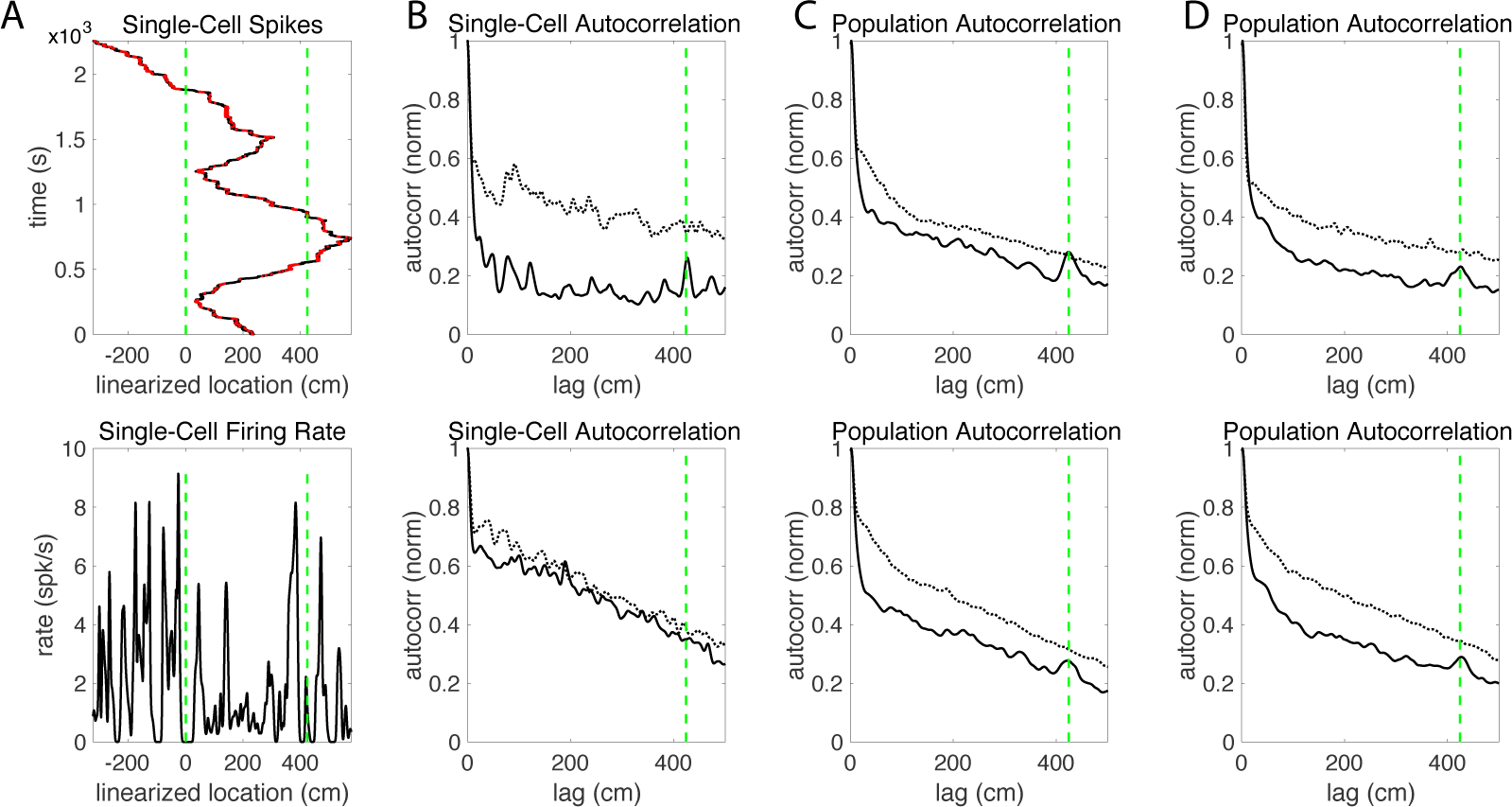
Analysis of the allocentric code for space. (A) Linearization of the circular track for an example cell. Dashed vertical lines mark one particular location on the circular track, which has a circumference of 424 cm. For the top plot, the rat’s linearized trajectory is shown in black, and the location of the rat dots indicate the time and linearized location of each spike. The bottom plot shows the corresponding firing rate on the linearized track. No consistent spatial structure is discernible from the firing rates of individual cells. (B) 1D autocorrelation of the firing rate for an example cell in Rat 1 (top) and Rat 2 (bottom). The autocorrelation of individual cells also does not show a consistent spatial structure and cannot be differentiated from the control (gray), which were generated by shuffling all spikes randomly across time. (C) 1D population autocorrelation, or sum of the 18 individual autocorrelations from Rat 1 (top) or 15 individual autocorrelations from Rat 2 (bottom) from the circular track in the light condition (*E*_2_). Both show a clear peak at the circumference of the track which clearly has a higher amplitude than any peak in the control. (D) Same as (C), but for the dark condition (*E*_3_). The peak at the circumference of the track is again clearly visible.

### Pairwise rate map correlations indicate a phase shift of grid cell activity between the open arena and the circular track environments

Our analyses using the population autocorrelation indicates that in this experiment, the spacing and orientation of the population lattice are largely preserved across environments *E*_1_, *E*_2_ and *E*_3_. However, the population autocorrelation cannot be used to determine whether there is a phase shift of the lattice between environments. Given that the circular track in *E*_2_ and *E*_3_ is the outer ring of the open arena *E*_1_, we next examine whether the hexagonal lattice of individual cells on the circular track is a circular slice of the same hexagonal lattice in the open arena.

To address this question, we characterize the similarity of the activity of individual grid cells across environments. Specifically, we extract the firing rates of each grid cell within the same ring of each environment (Fig. 8A). For each grid cell, we then compute the Pearson correlation coefficients for its extracted rate maps in *E*_1_ and *E*_2_, *E*_1_ and *E*_3_, and *E*_2_ and *E*_3_ (Fig. 8B-E, colored). As a control, we compare these same-cell correlations to the pairwise correlations of the extracted rate maps of difference cells (Fig. 8B-E, gray).

**Figure 8.**
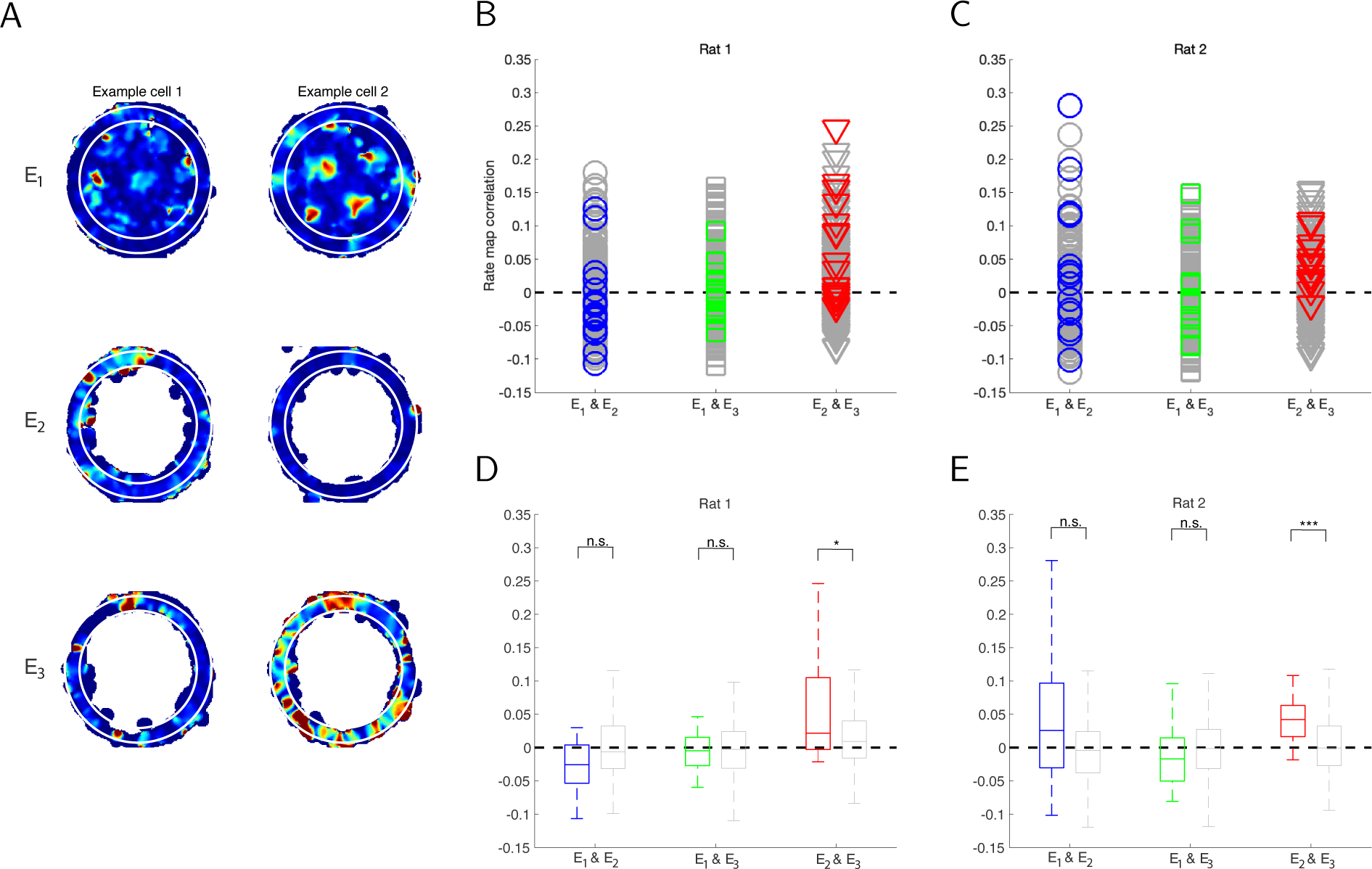
Spatial stability of firing fields across environments. (A) Examples of rate maps in each environment used for comparison. For each grid cell, we compute the pairwise correlations of its rate map in the two circular track environments (*E*_2_ and *E*_3_) and the corresponding slice of the open arena (*E*_1_, outlined by white circles). (B-C) Rate map correlations within the same cells (colored) and across cells (gray). (D-E) Box plot of the correlations shown in B-C, where the same-cell correlations are shown in color (left box plots) and the cross-cell correlations are shown in gray (right box plots). For both rats, there is a significant correlation between the rate maps on the circular track in the light and dark conditions (*E*_2_ and *E*_3_), but not between either circular track and the open arena. (For Rat 1, the p-values are 0.96 for *E*_1_ and *E*_2_, 0.48 for *E*_1_ and *E*_3_, and 0.02 for *E*_2_ and *E*_3_; for Rat 2, the p-values are 0.09 for *E*_1_ and *E*_2_, 0.77 for *E*_1_ and *E*_3_, and 0.001 for *E*_2_ and *E*_3_.) This indicates that there is some spatial stability in the rate maps of individual cells across conditions on the circular track, but the hexagonal lattice may have a phase shift between the open arena and circular track environments.

We find that for both Rat 1 and Rat 2, the extracted rate maps from the circular track in the light and dark conditions (*E*_2_ and *E*_3_) are significantly correlated, but the extracted rate maps from the open arena and either circular track environment (*E*_1_ and *E*_2_, *E*_1_ and *E*_3_) are not significantly correlated (Fig. 8D-E). Thus, in this experiment, the activity of individual grid cells is largely preserved in the same environment under different brightness conditions, but it is not preserved between the open arena and either circular track environment. This latter result is consistent with the reported observation that the grid cell activity on the circular track is not a circular slice of the activity in the open arena (Jacob et al., 2019).

If one were to consider only this pairwise correlation analysis, the lack of a significant correlation between the open arena and the circular track environments would appear to be a complete spatial remapping of the firing fields. When taken together with the population autocorrelation analyses, however, the most coherent explanation is that the hexagonal lattice of individual cells is maintained between environments, but there is a phase shift between the open arena and the circular track environments. Further analyses would be needed to determine if this phase shift is consistent across cells within a module, as would be predicted by attractor network theory.

## DISCUSSION

We demonstrate a novel statistical method for analyzing grid cell activity in undersampled environments, such as when the animal is constrained to a track. In these environments, the population autocorrelations reveal the underlying spatial structure of the grid cell activity, which cannot be discerned through the autocorrelations of individual grid cells. We first test the method through numerical simulations, showing that the method is robust to the spacing of the grid lattice, the width of the track, and realistic variations in the grid spacing and orientation among comodular grid cells. We then apply the method to experimental data. In Rat 1, the population autocorrelation clearly reveals a hexagonal lattice on the circular track in both light and dark conditions. The results in Rat 2 are less clear because the grid cells appear to be in two modules, and so there are fewer cells used in each population autocorrelation. Even in this case, however, a hexagonal lattice is revealed in the population autocorrelation in some of the circular track environments. In both cases, the grid spacing and orientation appear to be consistent between environments, though the phase appears to shift between the open arena and circular track.

### Comparison of the population autocorrelation to other methods

Autocorrelation is a powerful mathematical tool for finding repeating patterns. It has been widely used in data analysis and signal processing across disciplines. In neuroscience, it has become the standard approach for identifying grid cells in open-field environments. The usefulness of the autocorrelation is generally limited given insufficient or missing data. In the case of individual grid cell activity on a circular track, only some of the firing fields align with the track, leading to only a subset of the peaks appearing in the autocorrelation. The population autocorrelation successfully reveals the entire hexagonal pattern because the individual autocorrelations align with one another, and so the summation shows a more complete pattern than its individual components. Comodular grid cells are ideal for this approach because they differ only in phase, and the autocorrelation is independent of phase.This study is the first to apply the population autocorrelation to analyze grid cell data, though a similar approach using the population spectral density has been used to identify patterns in populations of place cells (Mulders et al., 2021).

Other techniques have been used to analyze grid cell data in undersampled environments. The Fourier transform has been used to determine that grid cell activity on a linear track is most likely a linear slice of an underlying 2D hexagonal lattice (Yoon et al., 2016). However, this approach does not extend naturally to nonlinear tracks, and we found in numerical simulations that there is no identifiable signature in the peaks of the Fourier transform given a circular slice of an underlying hexagonal lattice (data not shown). In contrast, the population autocorrelation can be used to analyze grid cell activity in any shape environment.

A second approach using methods from topology has also been used to conclude that grid cell activity in a pinwheel maze arises from an underlying hexagonal lattice (Gardner et al., 2022). While this approach would be effective in an environment of any shape, it requires simultaneous recordings from hundreds of grid cells, an order of magnitude larger than provided by most experimental studies. In contrast, we show here that the population autocorrelation can reveal an underlying pattern using only 18 grid cells in Rat 1, and to a lesser degree, 9 grid cells in Module 1 of Rat 2 and 5 grid cells in Module 2 of Rat 2. Additionally, the population autocorrelation is a direct extention of the individual autocorrelations that are the standard approach used by experimentalists to analyze grid cell activity. Thus, it is more easily understandable in the field of neuroscience than these two methods.

The population autocorrelation is limited in that recordings from an open-field environment must be used to identify comodular grid cells to be included in the summation. Because tetrodes tend to drift over time, this requires one to record grid cells in an open-field environment each day in addition to the recordings in the undersampled environment of interest. It is possible that the population autocorrelation approach is robust to the inclusion of relatively few non-grid cells, since any individual spatial structure that is not consistent across cells may be lost in the summation (e.g. Fig. 1D).

### Comparison to previous conclusions using this data set

Our main conclusion, that grid cell activity on the circular track is a slice of a hexagonal lattice, differs from the conclusion made in the original study presenting this data (Jacob et al., 2019). The key difference between our approaches is that they analyzed individual grid cell activity, and we analyzed the population activity. While it is important to study the neural behavior of individual cells, any underlying spatial structure of individual activity patterns is not generally observable when the animal is constrained to a circular track. This spatial structure only becomes apparent when one combines the activity patterns of a sufficient number of grid cells with different phases but approximately the same spacing and orientation. It is reasonable to consider the population code when determining how grid cells encode space, as downstream neurons integrate inputs from multiple grid cells.

In Jacob et al. (2019), the conclusion that individual grid cell activity is not a circular slice of a hexagonal lattice was not the focus of the study, and a thorough analysis establishing this conclusion was not presented. They based their conclusion on two observations: that the Fourier transform does not have the characteristic three peaks found in Yoon et al. (2016), and that there is a low correlation between the grid cell activity on the circular track and that in the open arena. As previously mentioned, the mathematical theory behind the three peaks of the Fourier transform applies only to a linear track, and there is no such signature of grid cells on a circular track. The latter observation is consistent with our own observation that the hexagonal lattice appears to differ in phase between the open arena and the circular track environments. A shifted lattice is not surprising, as the grid cell lattice often shifts and/or rotates between environments. The main conclusion of Jacob et al. (2019) is that most individual grid cells have a path integration code for space rather than an allocentric code. In particular, when they computed the Fourier transform of individual activity on the linearized track, they found that the largest peak differs from the circumference of the track, indicating that grid fields shift between laps. They interpreted this shift as evidence that grid cells have a path integration code for space on a circular track rather than an allocentric code, as this shift can be explained by an error in path integration but is not expected of a stable allocentric code. In contrast, we computed the autocorrelation of the population activity on the linearized track and found one clear peak at the circumference of the track, leading us to conclude that the population of grid cells has an allocentric code for space (Fig. 7C-D). The individual autocorrelations have several peaks, and we were unable to draw conclusions about a consistent shift in individual grid fields from our own analysis. Analysis of individual grid cells is difficult given this data set, as animals were permitted to turn around and run in either direction on the track. The animals completed between one and five laps, but in most sessions the animal completed fewer than two laps. Therefore, there is limited data to determine a consistent lap-by-lap shift of firing fields. The allocentric code we detected in the population autocorrelation is not surprising given the spatial structure of the population autocorrelation, especially in Rat 1. Stable firing fields are a necessary condition for the population autocorrelation to be useful, as the firing rates are averaged over the entire session.

### Implications for attractor network theory

Our results have important implications for the attractor network theory of grid cells, as the prior conclusion presented in Jacob et al. (2019) was a challenge to the theory. In the numerous attractor network models of grid cells, the hexagonal lattice arises from structured synaptic connections among grid cells, sometimes mediated by inhibitory cells. These models allow for deformations in the hexagonal lattice of individual cells given corresponding deformations in the external environment, as observed experimentally. However, they would not predict that grid cells have a fundamentally different code on a circular track as opposed to an open-field environment or linear track, and deformations would not be expected in this case since the circular track is simply the outer ring of the familiar open arena. The primary implication of this study is that attractor network theory can continue to be used to design models and interpret data of grid cell activity in any spatial environment. Our approach can also be used to further elucidate the properties of grid cells from recordings on tracks of any shape.

## METHODS

### Model simulations

In our simulations (Figs. 1-2), the circular arena has a radius of 75 cm, and the circular track is 15 cm wide with the same outer radius. The circular track was obtained by removing an inner circle of radius 60 cm from the arena, as was done in the original study (Jacob et al., 2019). Simulated grid cells have an idealistic firing rate map in which the hexagonal lattice of firing fields are constructed through a summation of three two-dimensional sinusoidal gratings at angles *θ, θ* + 60*^◦^* and *θ* + 120*^◦^*, where *θ* is the orientation of the lattice (Solstad et al., 2006). In the example used to generate Fig. 1, *θ* = 14.9*^◦^*, and the grid spacing is 45 cm for all simulated grid cells. The phase of each grid cell was randomly drawn from a uniform distribution, modeling an idealistic module of grid cells. Grid cell responses on the circular track are the corresponding slices of the hexagonal lattice in the circular arena. The shuffled rate maps in each environment were generated by shuffling the firing fields in that environment, with no overlapping fields allowed. To determine how the track itself contributes to the autocorrelation (Fig. 1F), we constructed a binary rate map by setting the rate on the track to 1 and the rate off the track to 0.

### Data analysis

We analyzed published data from a study in which rats explored both an open circular arena and a circular track in light and dark conditions (Jacob et al. (2019): https://zenodo.org/record/ 4662650#.YmwY_trMI2y). We include grid cells in our analysis that were recorded in all three environments. A total of 33 grid cells from two rats (18 in Rat 1 and 15 in Rat 2) are included in our analyses. The data from the other rats in the study do not contain sufficient cells that were recorded in all three environments and have a hexagonal firing pattern in the open arena, and so we analyzed data only from grid cells in Rats 1 and 2.

### Average firing rate

The average firing rate of a given cell at any position **x** is given by

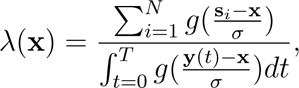

where *g*(*·*) is a smoothing Gaussian kernel with the smoothing factor *σ* = 3 cm, *N* is the total number of spikes of the cell within the recording session, **s***_i_* is the rat’s location at the time of the *i*-th spike, **y**(*t*) is the rat’s location at time *t*, and *T* is the total time of the recording session (Fyhn et al., 2004; Hafting et al., 2005). The rate at any unvisited location was set to 0, where a location **x** is considered unvisited when it is more than 3 cm away from all recorded animal positions, i.e., min*_t_ |***x − y**(*t*)|*>* 3. We then binned the rate maps, using a resolution of 1 cm × 1 cm for both rate maps and their 2D autocorrelations.

### Gridness score

Gridness scores were calculated from the 2D autocorrelation (either of an individual rate map or the population autocorrelation) using a similar approach to that of several prior studies (Hafting et al., 2005; Brandon et al., 2011; Savelli et al., 2017). We first extracted the annulus containing the innermost ring of peaks from the 2D autocorrelation, excluding the central peak. To determine this annulus algorithmically, we computed *M* (*r*), the mean autocorrelation along concentric circles or radius *r* centered at the origin. We observed that for autocorrelations with a clear hexagonal structure, *M* (*r*) decreases from the origin until reaching the first ring of peaks, at which point it increases until reaching a maximum value at the center of these peaks. Accordingly, we set *r*_min_ and *r*_max_ to be the radii at which *M* (*r*) is minimized and maximized, respectively. We then defined the extracted annulus to have an inner radius of *r*_min_ and an outer radius of *r*_max_ + (*r*_max_ *− r*_min_). See Supp. Figs. S1 and S8 for examples of autocorrelations with the circles of radius *r*_min_ and *r*_max_ depicted in white.

For cases in which the function *M* (*r*) does not have clear minimum and maximum values, which occurs when the autocorrelation lacks spatial structure, we set *r*_min_ = 25 cm and *r*_max_ = 40 cm. The qualitative results are robust to the choice of *r*_min_ and *r*_max_ in these cases.

Next, we computed the rotational autocorrelation of this extracted annulus. Given a perfect hexagonal lattice, the highest values of the rotational autocorrelation should be at 60*^◦^* and 120*^◦^*, while the lowest values should be at 30*^◦^,* 90*^◦^* and 150*^◦^*. The gridness score is the difference between the lowest value observed at 60*^◦^* or 120*^◦^* rotation and the highest value observed at 30*^◦^,* 90*^◦^*, or 150*^◦^* rotation.

### Inferring grid lattice features

A hexagonal lattice has four defining features: its spacing, orientation, and 2D phase. While the phase does not affect the autocorrelation, the spacing and orientation can be determined from the innermost ring of six peaks in the 2D autocorrelation. Since the autocorrelation has a 180*^◦^* rotational symmetry, any three peaks on a semi-annulus suffice. Beginning with the extracted annulus used to compute the gridness score, we located the three peaks in the upper annulus (from 0*^◦^* to 180*^◦^* from the positive *x*-axis). We did this by first locating the position of the maximum autocorrelation value in the upper annulus. We then excluded the segment of the annulus spanning 35*^◦^* from that point and located the position of the maximum autocorrelation from the remaining upper annulus. We then repeated this process once more to locate the third peak. See Supp. Figs. S1 and S8 for examples, where the three peaks are indicated by white crosses.

After locating the three peaks, we inferred the grid spacing as the average radial distance to the three peaks, and we inferred the three orientation angles as the angles from each of the three peaks to the positive *x*-axis. The three orientation angles should be roughly 60*^◦^* apart for a hexagonal lattice. These inferred grid features are shown in Figs. 4 and 5.

### Hierarchical clustering in Rat 2

To partition the grid cells in Rat 2 into two modules, we applied hierarchical clustering based on the inferred spacing and three orientation angles for each grid cell. The *z*-scores of the inferred spacing and orientations were used as the features, and the distance between clusters was defined as the average Euclidean distance between all pairs in the two clusters.

### 2D shuffle controls

We generated controls by shuffling the firing fields of the rate map for each cell such that no two fields overlap in the shuffled rate map. We define firing fields by first setting a threshold, and then finding connected pixels above the threshold using the MATLAB function bwconncomp. In order to preserve the size of the fields as much as possible, we start from zero threshold. We sort the connected components based on their areas, and randomly place them in the environment they are extracted without overlapping. The largest component is placed first, and the placement is in order of decreasing area. The zero threshold may not work in all cases since some grid cells fired outside their firing fields and had non-zero activity over a large part of the environment. In that case, random placement of one or a few large fields may render the subsequent placement of a field impossible. To avoid having an oversized field covering too much area outside the actual field, we set a constraint to the field size. Based on the rate maps, the maximum field sizes in the circular arena and track are constrained to 1000 and 500 cm^2^. When any field size exceeds the maximum constraint, or the subsequent placement of a field becomes impossible, the threshold for field detection is raised by 1% of the maximum value of the rate map. This procedure is repeated until all the detected fields are placed in the environment they are detected. To preserve the firing statistics of the rate map, the pixels with subthreshold activity are randomly assigned to the control rate map with zero activity.

Across all cells in all environments, the thresholds for field detection range from 0% to 15% of the maximum value of the rate map, with an average of 3.98%.

### Linearized activity

We linearized the path of the rat according to the track positions (Jacob et al., 2019). In particular, the position of the animal on the 1D linearized track is the total angular distance traveled by the rat from its starting position, where the distance is positive when the animal moves clockwise and negative when the animal moves counterclockwise. Spikes and timestamps are sorted into bins of 1 cm. The average rate at any bin was calculated by the number of spikes divided by the number of timestamps in that bin, and then divided by the time step. Controls are generated by shuffling all spikes randomly across time.

### Pairwise correlation of rate maps

For each cell, we extract an annulus of inner and outer radii of 120 cm and 150 cm, respectively, from the rate maps in all three environments. We then normalize the rate maps by subtracting the the average rate across visited locations on the annulus. The firing rate at unvisited locations on the normalized rate map is set to 0. This step is taken to eliminate potential bias due to different degrees of environment exploration across environments and across sessions. The rate profile of each extracted normalized rate map is linearized into a 1D vector. For each cell, the Pearson correlation coefficient is used to define the correlation of normalized rate maps among environments. Cross-cell correlations are used for controls, in which we compute the Pearson correlation coefficients of the normalized rate maps of different cells. The right-tailed Mann-Whitney *U* test (Nieuwenhuis et al., 2011) is used to determine the significance of the same-cell correlations across environments.

## SUPPLEMENTAL INFORMATION

Supplemental Information includes 11 figures and can be found with this article online.

## ACKNOWLEDGMENTS

This work was supported through the BRAIN Initiative and the Collaborative Research in Computational Neuroscience (CRCNS) program, grant 1R01MH118926-01. We thank Pierre-Yves Jacob and Francesca Sargolini for sharing their data and discussions. We also thank James Knierim, Noah Cowan, Kechen Zhang, Francesco Savelli, Gorkem Secer, Manu Madhav, Ravikrishnan Jayakumar, and Bharath Krishnan for their helpful feedback and discussions. Computational resources for this research were provided by SMU’s Center for Research Computing.

## AUTHOR CONTRIBUTIONS

Conceptualization, M.Y.Y. and K.H.; Methodology, M.Y.Y. and K.H.; Data Analysis, M.Y.Y. and S.W.; Model Simulation, M.Y.Y.; Writing, M.Y.Y. and K.H..

## Supplementary Figures

**Figure S1.**
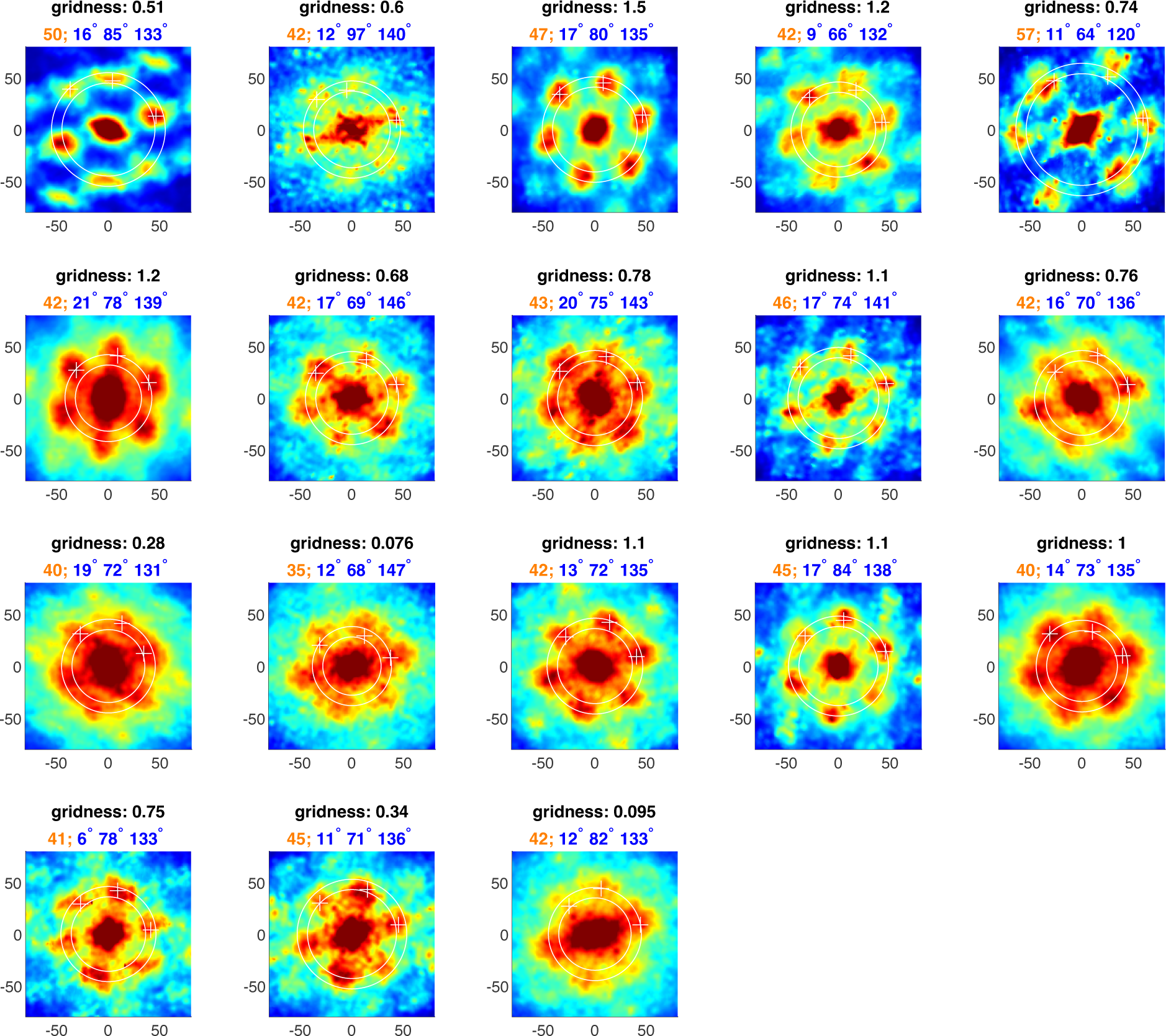
Autocorrelations of all 18 grid cells in Rat 1 in the open arena. (Environment *E*_1_). The inferred spacing and orientations are labeled with white circle and white pluses, respectively. Those values (black), together with the gridness score (blue), are shown for each cells.

**Figure S2.**
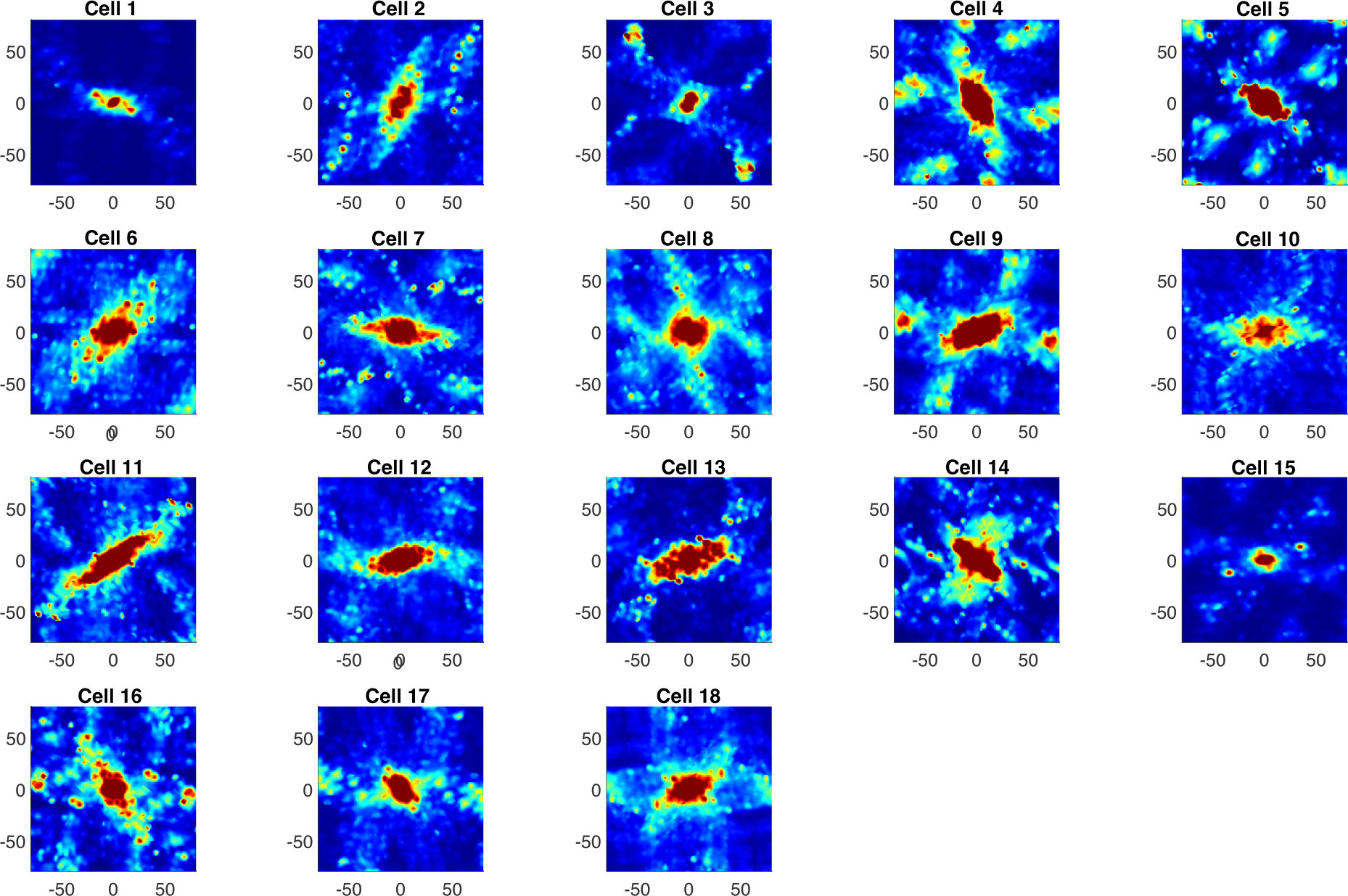
Autocorrelations of all 18 grid cells in Rat 1 on the circular track in the light condition. (Environment *E*_2_). A few cells have peaks that could be part of an underlying hexagonal lattice (e.g. Cells 5, 9, 15 and 17). However, the spatial structure of even these cells is unclear, and overall the individual autocorrelations do not clearly indicate a particular consistent spatial structure in the grid cell firing fields.

**Figure S3.**
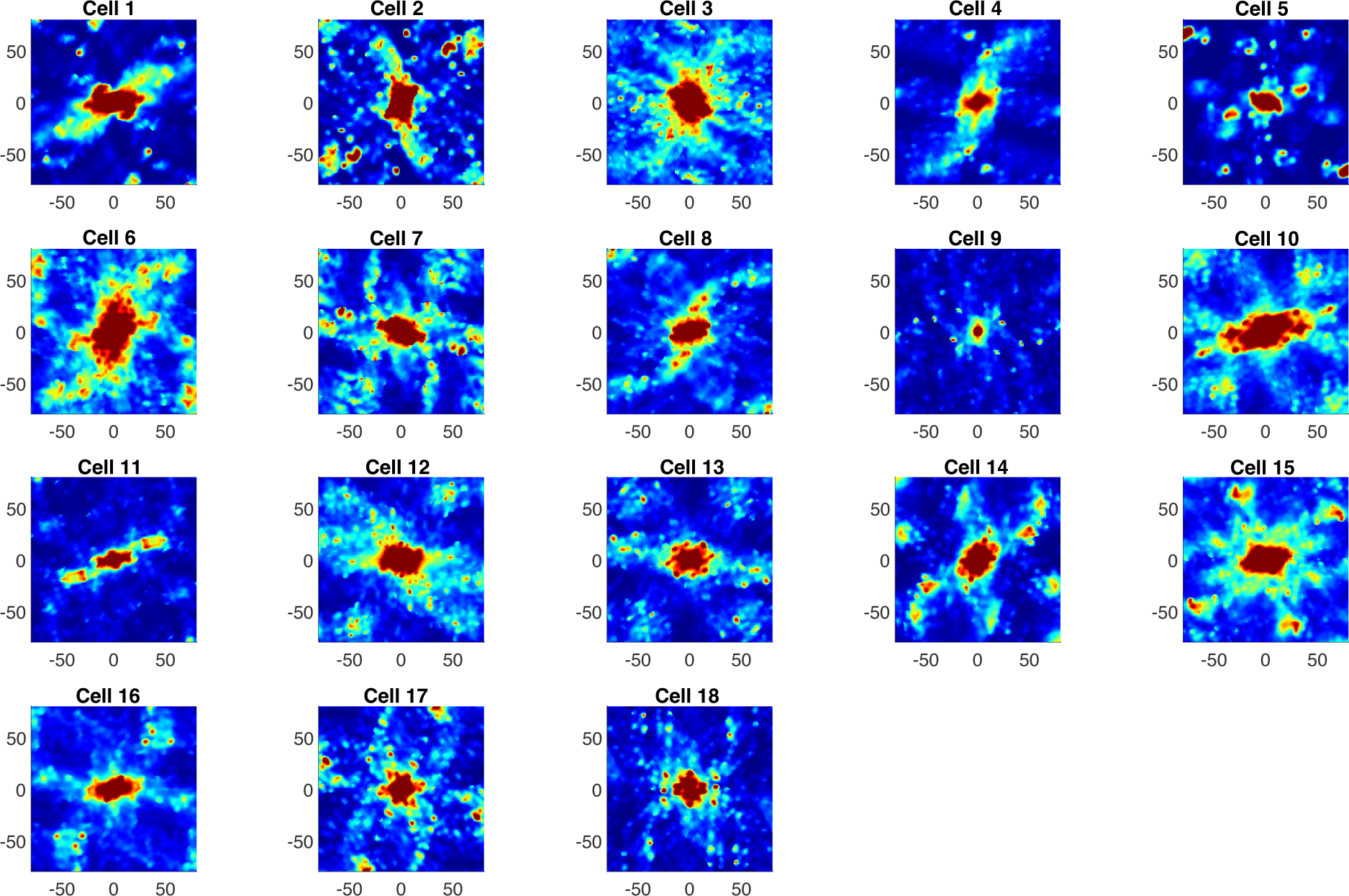
Autocorrelations of all 18 grid cells in Rat 1 on the circular track in the dark condition. (Environment *E*_3_). Like in *E*_1_, a few cells have peaks that could be part of an underlying lattice (e.g. Cells 13-16). However, the overall spatial structure is again unclear from these individual autocorrelations.

**Figure S4.**
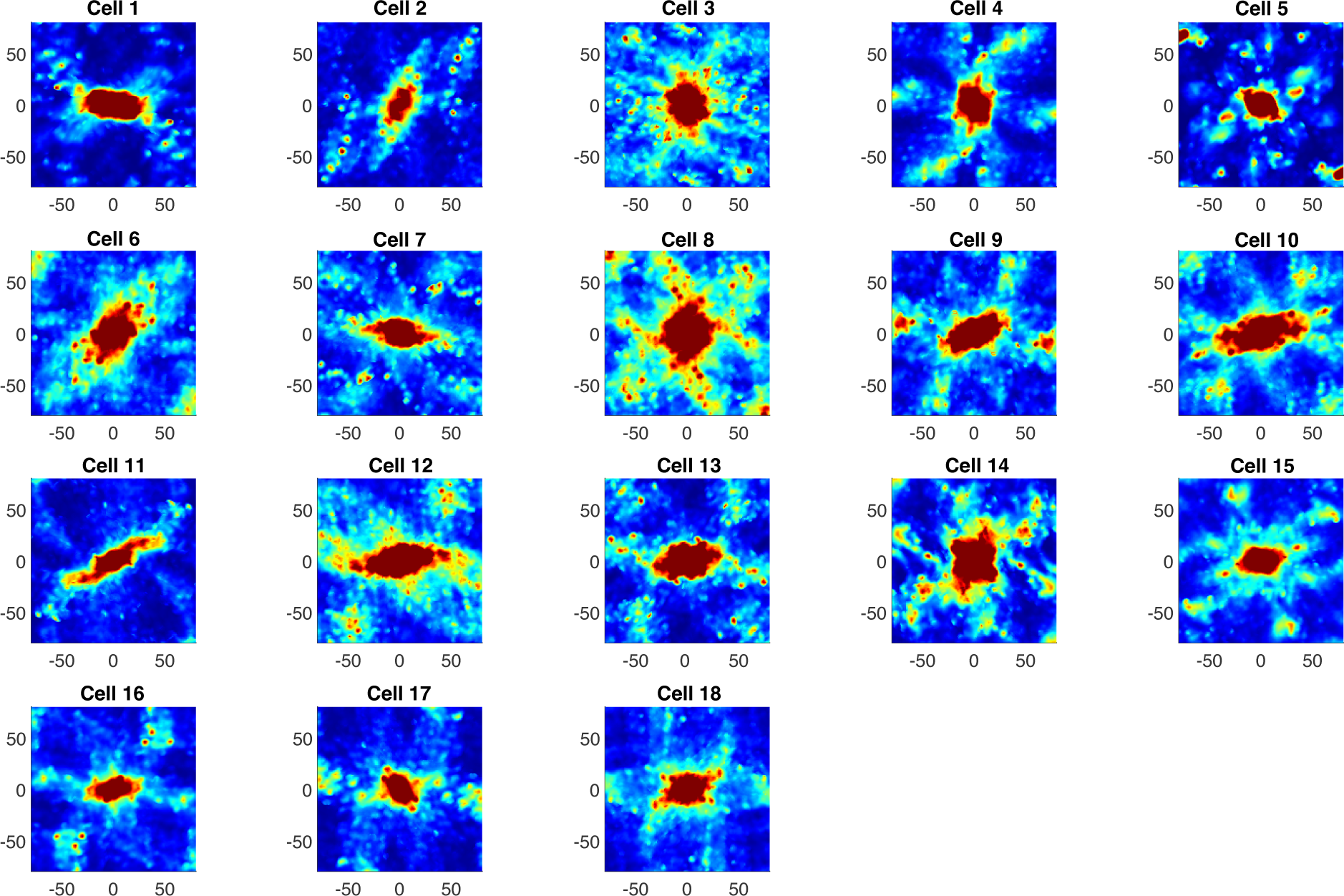
Combined autocorrelations of all 18 grid cells in Rat 1 on the circular track in both conditions. The combined autocorrelation for each grid cell is the sum of the autocorrelations in *E*_1_ and *E*_2_. While there is still no consistent spatial structure, a few cells show the ring of 6 evenly spaced peaks characteristic of a hexagonal lattice. Note that the autocorrelation of Cell 8 resembles the hexagonal structure, while only two peaks appear in *E*_2_ and four in *E*_3_.

**Figure S5.**
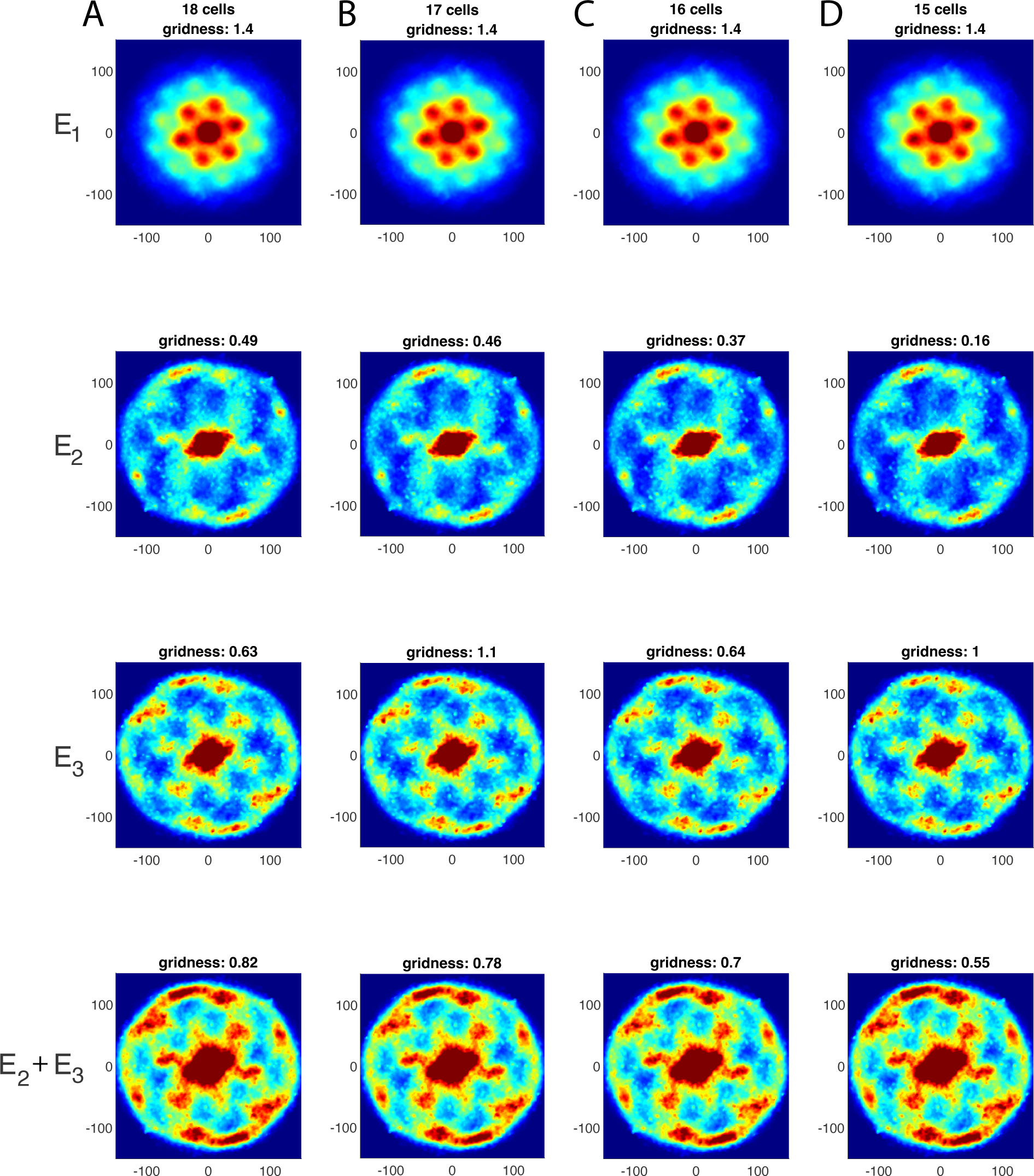
Robustness of results in Rat 1 to removing structured grid cells. To verify that the hexagonal structure in the population autocorrelation is a population effect that is not due to a few dominant grid cells, we recomputed the population autocorrelation while leaving out the grid cells with the clearest hexagonal structure in their individual autocorrelations in the combined environments (*E*_2_ +*E*_3_, shown Supp. Fig. S4). (A) Population autocorrelation using all 18 grid cells. (B-D) Population autocorrelation using all cells except Cell 14 (B), Cells 14 and 8 (C), and Cells 14, 8 and 16 (D). The hexagonal structure is robust to removing these cells, particularly in Environments *E*_2_ and *E*_2_ + *E*_3_.

**Figure S6.**
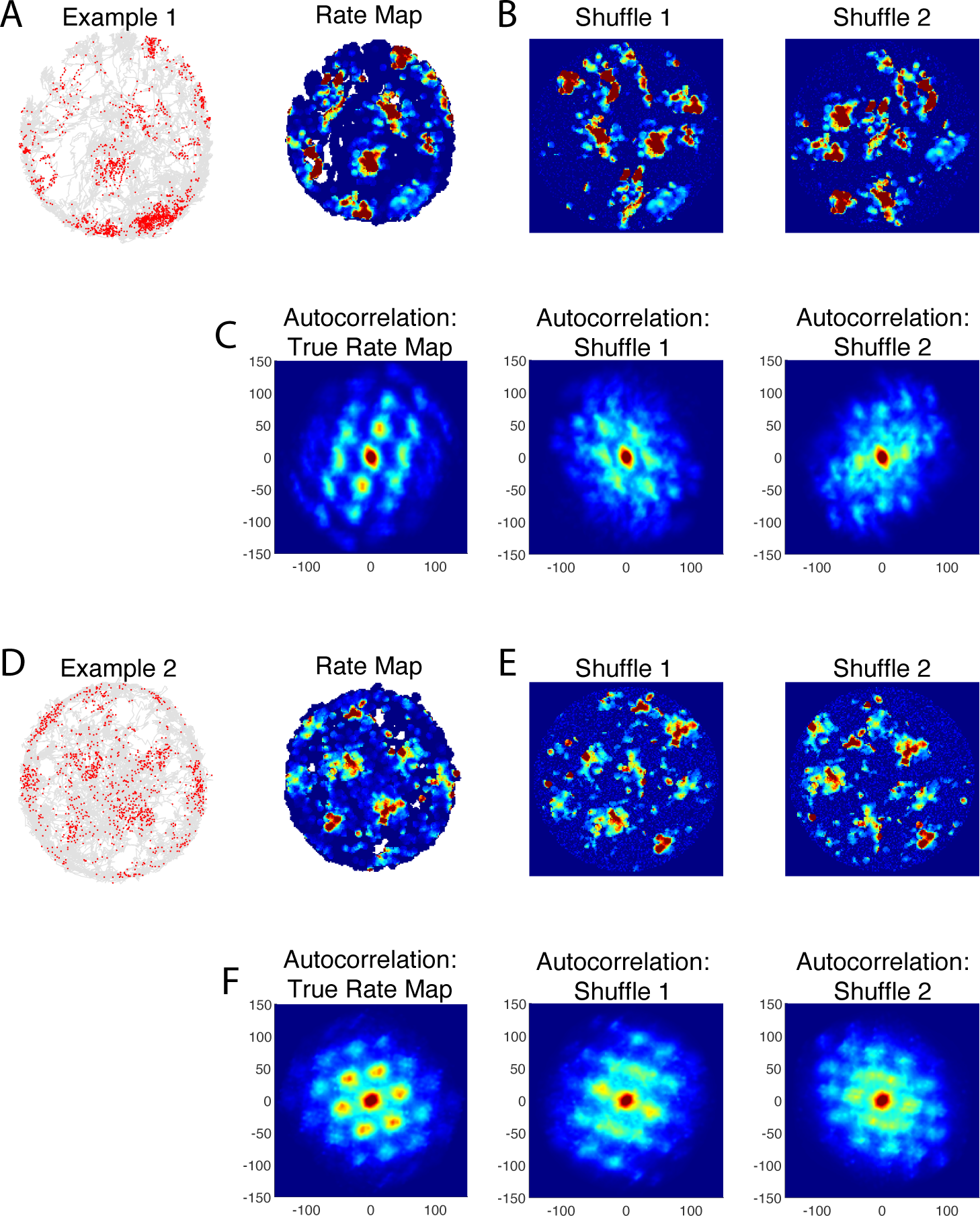
Examples of shuffle controls in the open arena. (Environment *E*_1_). (A) Data from an example cell in Rat 1. On the left, each red dot shows the location of the rat when this grid cell spiked, and the gray shows the rat’s trajectory. The corresponding rate map is shown on the right. (B) Two example shuffles of the firing fields in the rate map. Note that the firing fields are shuffled, not the individual spikes. Thus the non-spatial firing rate statistics are preserved in the shuffles. (C) Autocorrelations of the true rate map, Shuffle 1 and Shuffle 2. (D-F) Same as A-C, but for a second example cell. As expected, the hexagonal pattern is much clearer in the autocorrelation of the true rate map than for the shuffles.

**Figure S7.**
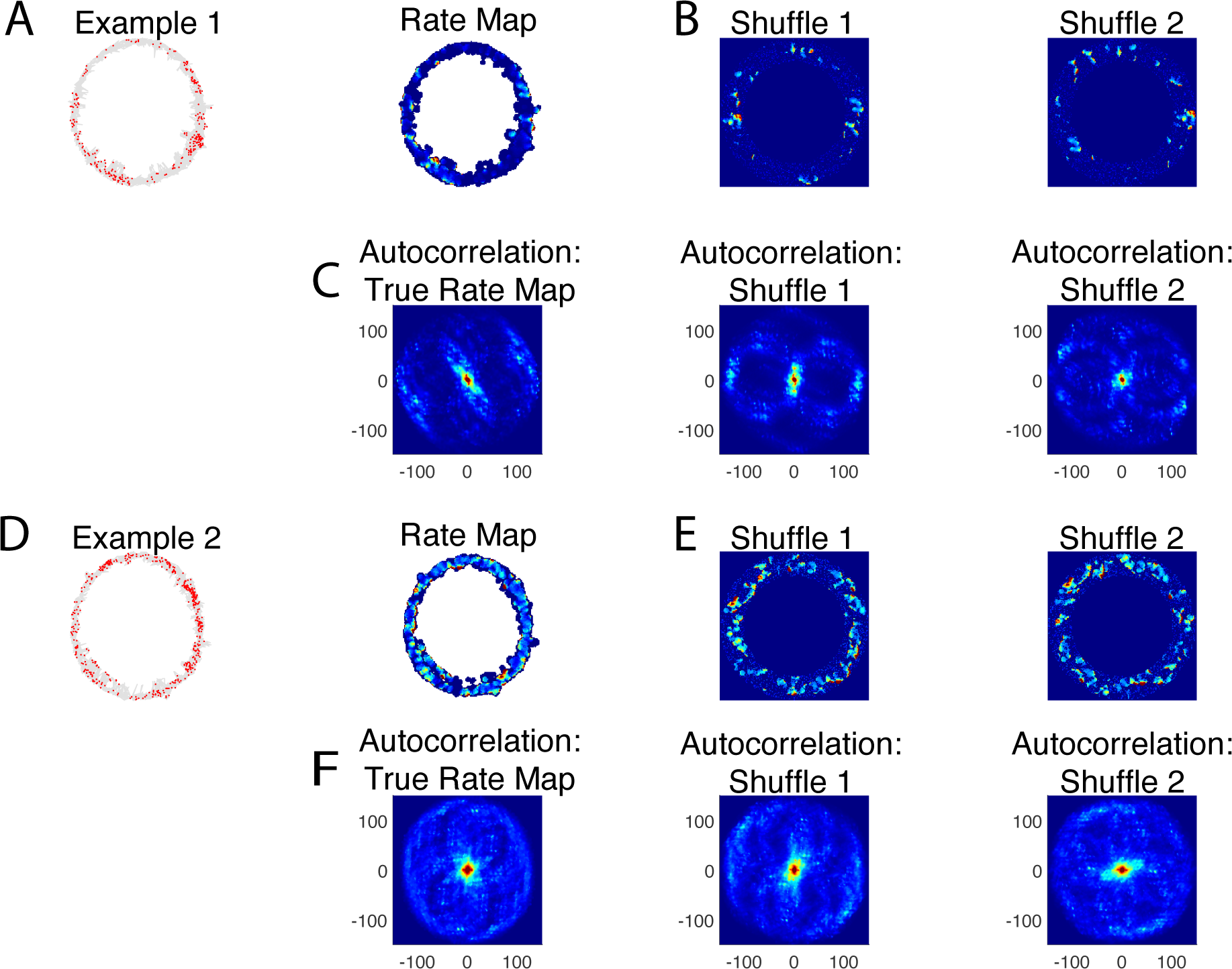
Examples of shuffle controls on the circular track. (Environment *E*_3_). (A) Data from an example cell in Rat 1. On the left, each red dot shows the location of the rat when this grid cell spiked, and the gray shows the rat’s trajectory. The corresponding rate map is shown on the right. (B) Two examples shuffles of the firing fields in the rate map. As done for the open field environment, firing fields are kept intact so that the non-spatial statistics are preserved in the shuffles. (C) Autocorrelations of the true rate map, Shuffle 1 and Shuffle 2. (D-F) Same as A-C, but for a second example cell. The autocorrelations of individual cells are usually not informative about the underlying spatial firing pattern due to the sparsity of the data on the circular track.

**Figure S8.**
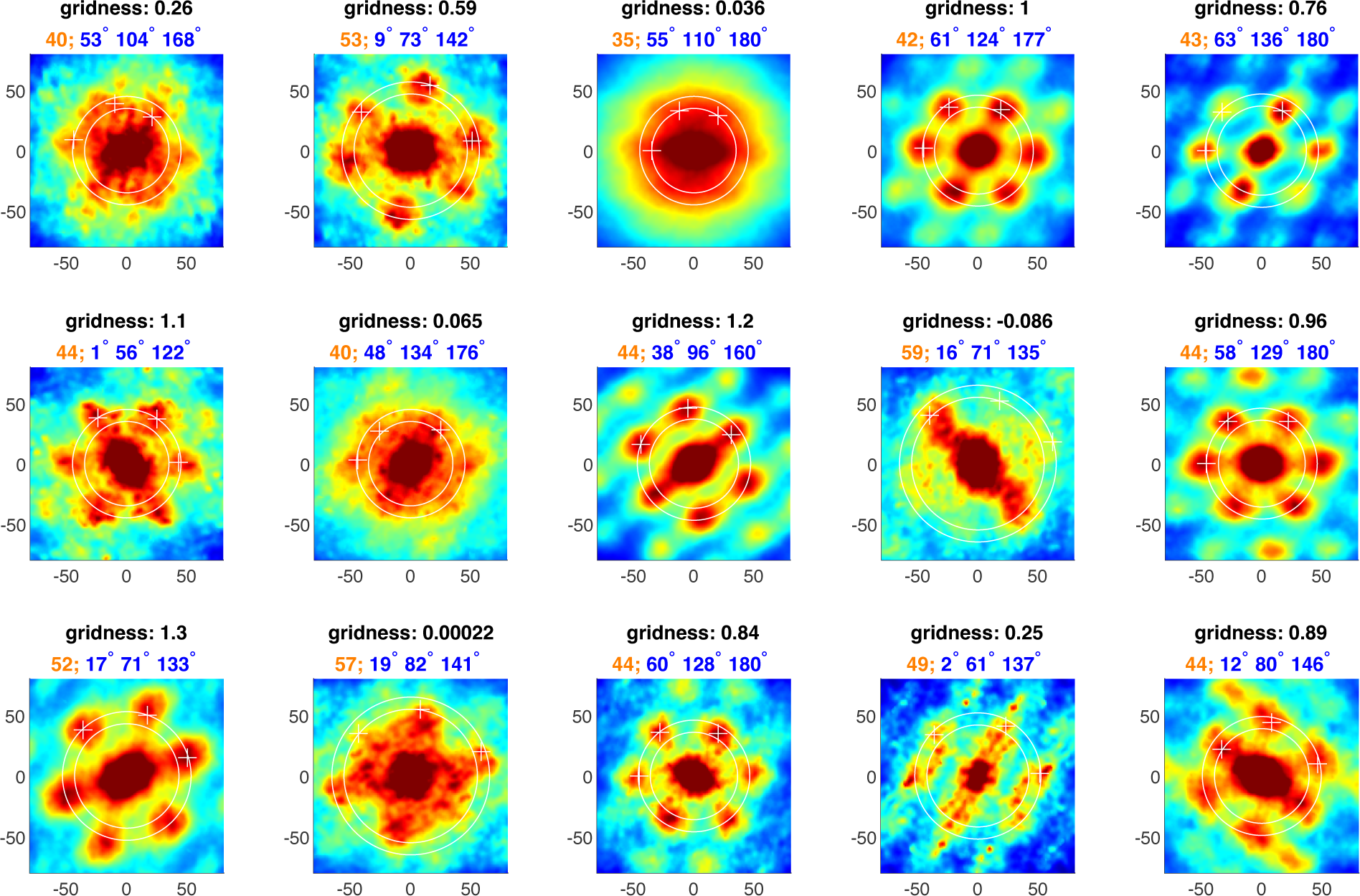
Autocorrelations of all 15 grid cells in Rat 2 in 2D arena.

**Figure S9.**
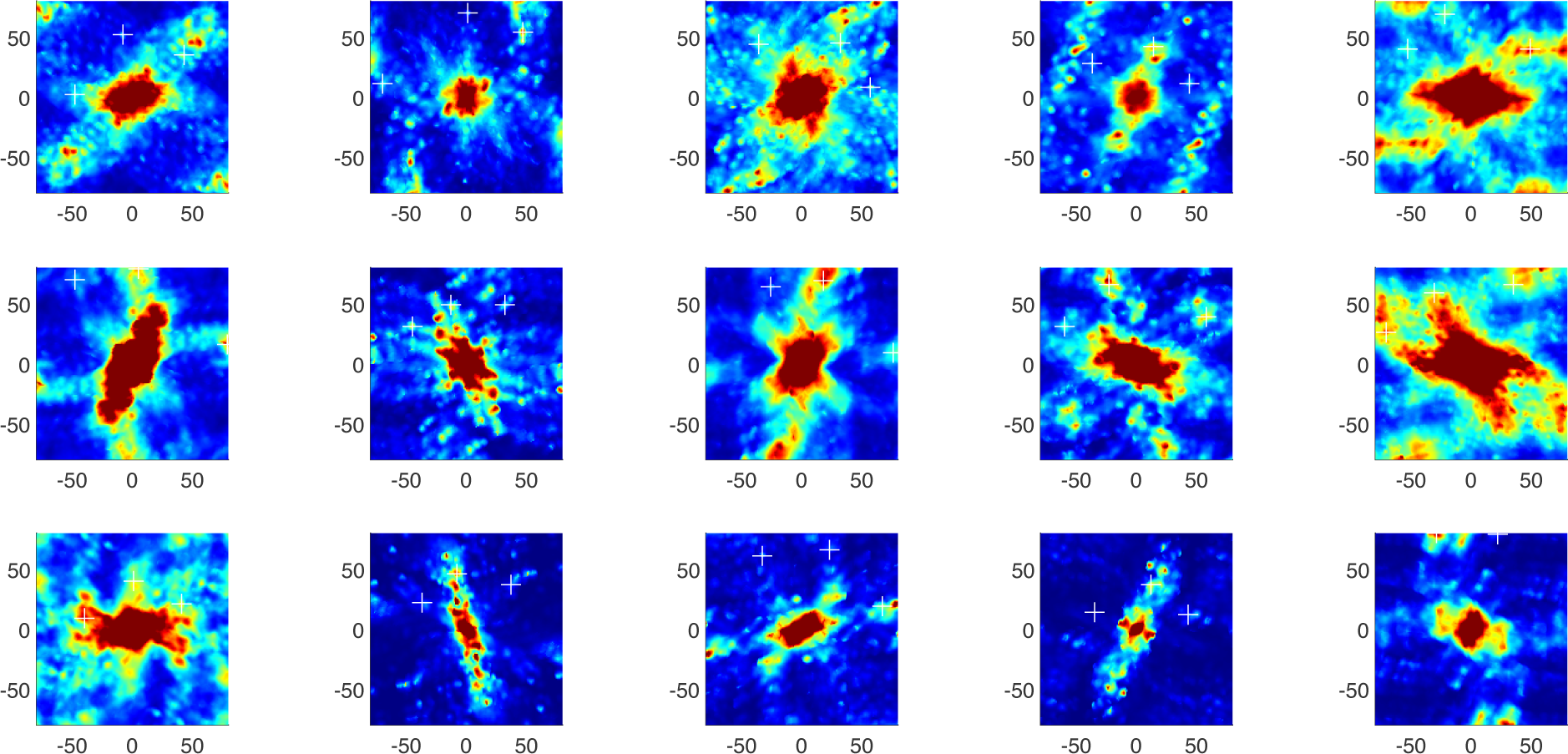
Autocorrelations of all 15 grid cells in Rat 2 on the light annlus.

**Figure S10.**
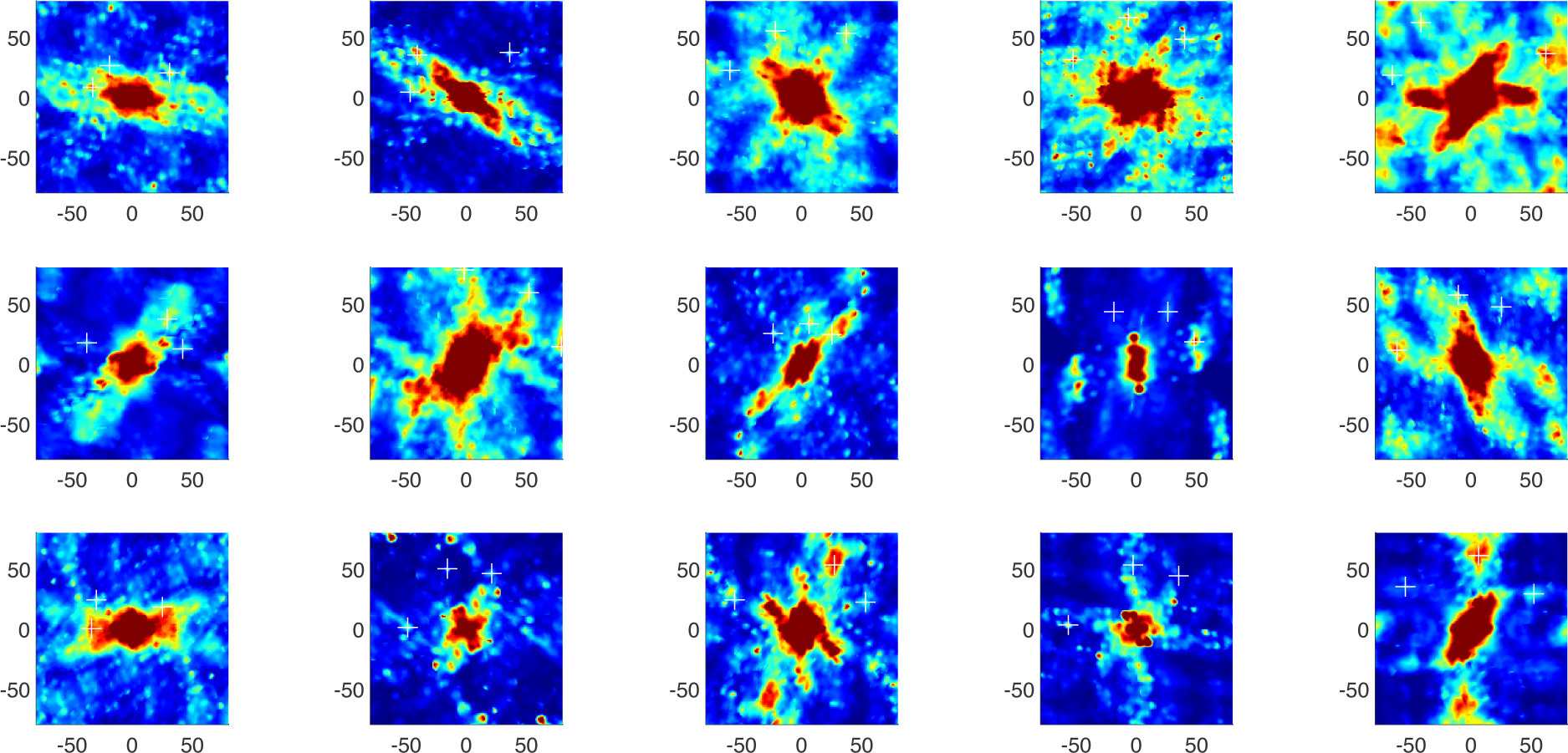
Autocorrelations of all 15 grid cells in Rat 2 on the dark annulus.

**Figure S11.**
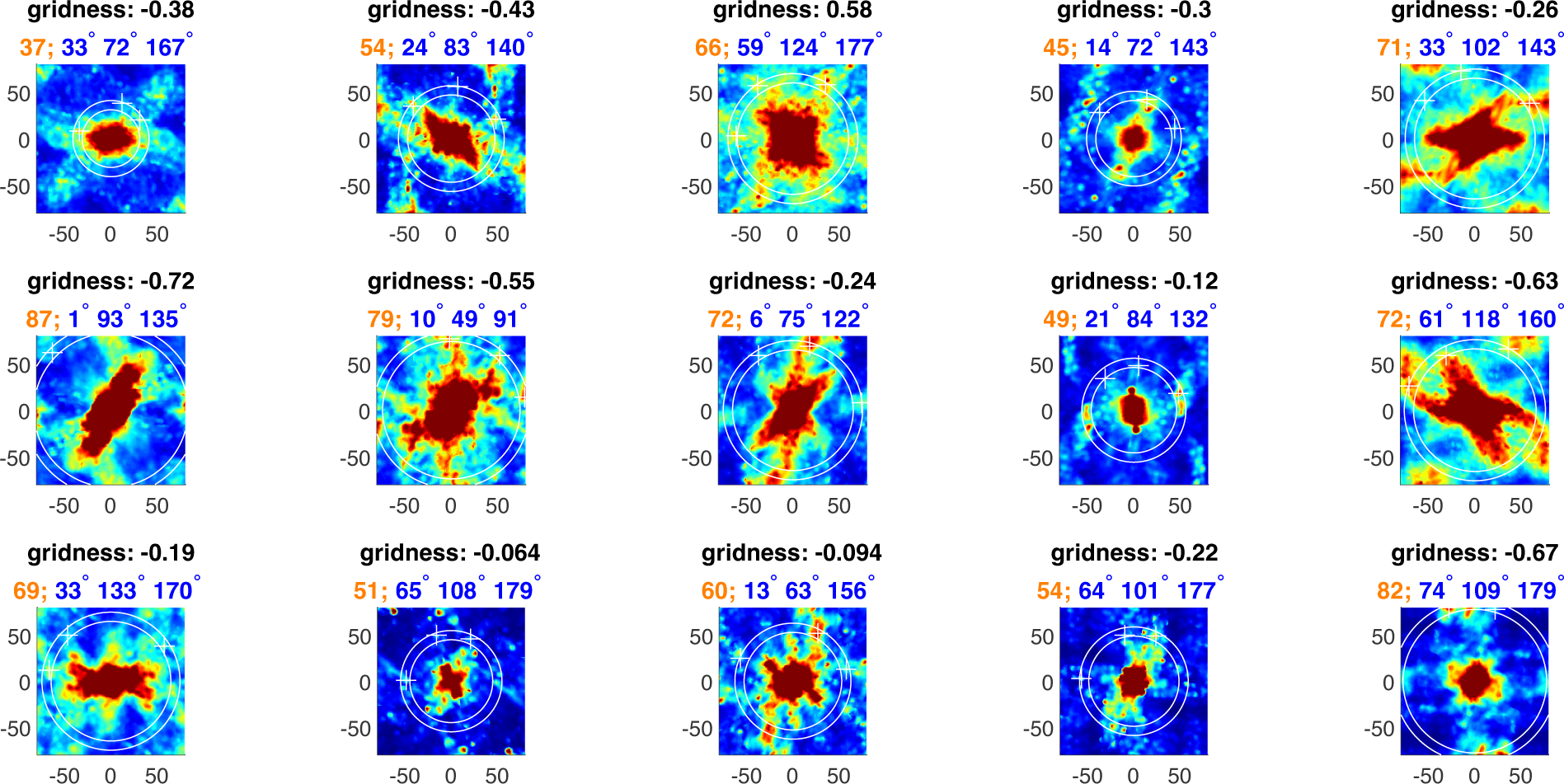
Autocorrelations of all 15 grid cells in Rat 2 on the light and dark annuli combined.

